# Living Photovoltaics based on Recombinant Expression of MtrA Decaheme in Photosynthetic Bacteria

**DOI:** 10.1101/2023.02.28.530417

**Authors:** Melania Reggente, Nils Schurgers, Mohammed Mouhib, Sara Politi, Alessandra Antonucci, Ardemis A. Boghossian

**Affiliations:** École Polytechnique Fédérale de Lausanne (EPFL), 1015-Lausanne, Switzerland; Dipartimento di Scienze e Tecnologie Chimiche, Università degli Studi di Roma “Tor Vergata”, Via della Ricerca Scientifica, 00133, Rome, Italy; Institute of Biology III, University of Freiburg, 79104 Freiburg, Germany

**Keywords:** bioengineering, cytochrome, MtrCAB, extracellular electron transfer (EET), direct electron transfer (DET), exoelectrogenicity, microbial fuel cell (MFC), biological photovoltaic or biophotovoltaic (bioPV, BPV), photosynthesis

## Abstract

At the center of microbial bioelectricity applications lies the critical need to express foreign heme proteins that are capable of redirecting the electron flux of the cell’s metabolism. This study presents bioengineered *Synechocystis* sp. PCC 6803 cells capable of increased electrogenicity through the introduction of a genetic construct for cytochrome expression. We could demonstrate the functional expression of the periplasmic MtrA decaheme c-type cytochrome from *Shewanella oneidensis*, a dissimilatory metal-reducing exoelectrogen, inside *Synechocystis*. Protein expression was verified through western-blotting and immunostaining, and oxygen evolution, optical density, and absorption measurements confirm sustained cell activity and viability under the tested expression conditions. Furthermore, the bioengineered cells show enhanced mediated exoelectrogenicity, as confirmed through a colorimetric iron assay and electrochemical measurements. Compared to wildtype cells on graphite electrodes, the bioengineered cells show a 2-fold increase in light-dependent, extracellular electron transfer, achieving photocurrent densities of 4 μA/cm^2^ under white light illumination of ∼500 μmol m^-2^s^-1^. The increased capacitance obtained under illumination and suppressed photocurrents in the presence of the photosynthetic inhibitor, 3-(3,4-dichlorophenyl)-1,1-dimethylurea (DCMU) suggest increased extraction of photosynthetically derived electrons from the recombinant cells. The improved bioelectricity transport across the outer membranes, as achieved through the heterologous heme expression inside cyanobacteria, presents new opportunities for re-wiring the metabolisms of light-harvesting microbes.

## Introduction

Living photovoltaics harness the metabolisms of live cells to convert solar energy into electricity. The incorporation of photosynthetic microbes in these devices presents significant advantages that reach beyond the limits of conventional inorganic devices. For example, photosynthetic microbes benefit from adaptive light-harvesting and dynamic self-repair processes that impart biological devices with fault-tolerance [1]. In addition, compared to inorganic photovoltaics that release substantial quantities of CO_2_ during processing [2] photosynthetic microbes actively sequester CO_2_ during their growth and doubling cycles, contributing to an overall negative carbon footprint. When coupled with their ability to replicate, their reliance on abundant and accessible raw materials further gives rise to a self-sustaining technology for covering spacious surfaces that are ideally suited for solar energy harvesting [3].

Despite their advantages, living photovoltaics suffer from low conversion efficiencies that are largely limited by inefficient extracellular electron transfer (EET) across the outer membranes of the living cells [4]. Though previous studies have reported low photocurrent production from wildtype photosynthetic microbes such as cyanobacteria and algae [4-7], the EET mechanism remains largely elusive and poorly understood [8]. By contrast, non-photosynthetic, metal-reducing microbes such as *Geobacter metallireducens* and *Shewanella oneidensis* MR-1 have evolved innate EET pathways that render them ideal for bioelectricity applications [9]. The most characterized EET pathway in *S. oneidensis* MR-1 involves the MtrCAB complex, [10,11] whose crystal structure has only recently been solved [12]. (**Figure 1**). Based on the current understanding of this pathway, electrons extracted from the oxidation of a substrate are ultimately transferred to the periplasmic decaheme cytochrome MtrA during anaerobic respiration. Subsequent electron transfer to the outer membrane decaheme cytochrome MtrC is facilitated by the outer membrane β-barrel protein MtrB, which places MtrA in direct contact with MtrC [13,14]. MtrC is re-oxidized by reducing terminal electron acceptors, such as electrodes, *via* direct or mediated electron transfer [14]. The former requires physical contact between the electrode and the outer membrane redox proteins, while the latter involves the use of a soluble electron shuttle that can be reduced by the outer membrane cytochromes and carry the electrons to the electrode in this reduced state.

**Figure 1.**
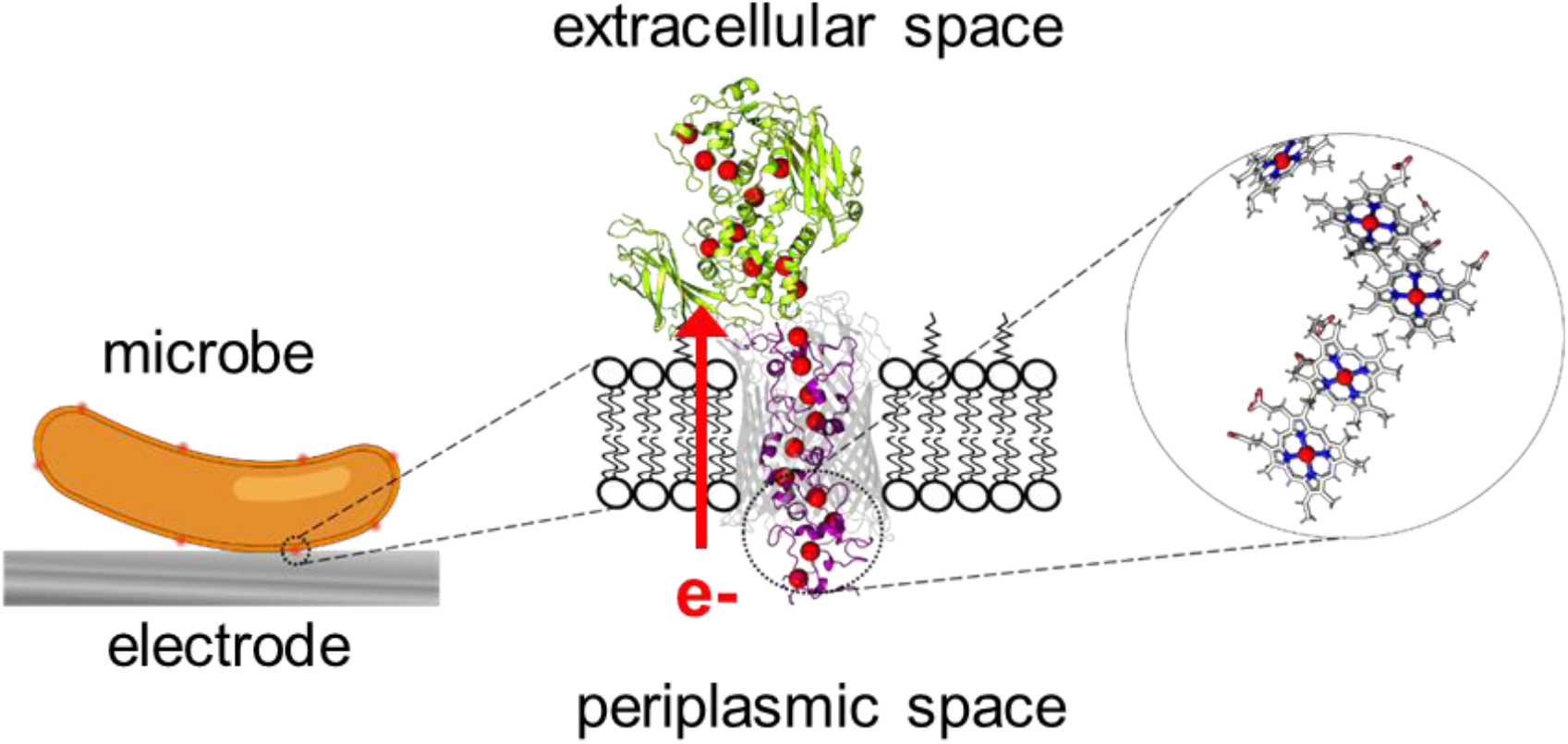
The MtrCAB Complex in the Outer Membrane of *S. oneidensis* MR-1. (Left) Metal-reducing microbes such as *Shewanella oneidensis MR-1*, shown in orange, interface with solid electrodes, shown in gray in microbial fuel cells. (Center) In the outer membrane, the MtrCAB pathway enables direct electron transfer to the electrode. Electrons are translocated from MtrA in the periplasm to MtrC in the extracellular space. MtrA is shown in magenta, MtrB in gray, and MtrC in green. The bilayer lipid membrane is designated in black. The heme centers are shown as red spheres. (Right) The closely stacked heme cofactors in MtrA facilitate electron transfer within the protein and across the membrane. (PDB: 6r2q [12])

Although metal-reducing microbes show increased EET efficiencies compared to their photosynthetic counterparts, they rely on the oxidation of specific and limited organic substrates for bioelectricity generation [15,4]. They further lack the CO_2_-sequestering metabolism found in photosynthetic organisms. Importantly, they also lack the light-harvesting machinery needed for water-splitting, an otherwise thermodynamically unfavorable reaction. This water-splitting reaction is key for the next generation of renewable energy technologies that rely on hydrogen production or even direct bioelectricity generation from ubiquitously abundant and accessible substrates like water.

To this end, synthetic biology offers a promising prospect for re-purposing a photosynthetic microbe’s metabolism for solar bioelectricity generation. Synthetic biology has already brought significant advancements to the realms of health [16], environment [17], and energy [18] though it has played a limited role in bioelectricity applications. Examples of bioengineering on this front are scarce, and have been largely restricted to the few non-photosynthetic microbes that show innate EET. For example, *S. oneidensis* MR-1 has been bioengineered to improve biofilm formation, [19], broaden and improve the utilization of various substrates [20, 21], and tune the expression of endogenous metabolites [22-24] and electron conduits [25, 26] to ultimately improve overall fuel cell performance [27]. Recent efforts have also focused on introducing EET pathways in non-exoelectrogenic microbes through the heterologous expression of EET conduits or outer membrane redox proteins. [28, 29]. For example, *E. coli* strains have been genetically modified to express the complete MtrCAB pathway, resulting in enhanced bioelectricity generation from an otherwise non-exoelectrogenic microbe [28, 30-35]. In addition, the autotrophic strain, *Synechococcus elongatus* PCC 7942, was also genetically engineered to express the outer membrane cytochrome S (OmcS) [29, 36]. Other bioengineering approaches to improve bioelectricity generation have focused on eliminating competitive electron pathways by suppressing the expression of terminal oxidases in cyanobacterial strain, *Synechocystis sp*. PCC 6803 [37] or on endogenously producing phenazines as diffusional mediators [38]. These efforts have all proven vital for improving the fuel cell efficiencies of natural exoelectrogens as well as extending the fuel cell applications to a collection of otherwise non-exoelectrogenic microbes with augmented and complementary metabolisms. However, despite these advancements, the bioengineering of defined EET pathways in photosynthetic microbes like cyanobacteria remains largely hindered by the need to express foreign cytochromes inside the cell which, to our knowledge, has yet to be achieved with *Synechocystis*.

The goal of the study herein is to bioengineer a living photovoltaic through the recombinant expression of a foreign cytochrome inside cyanobacteria. This work establishes a genetic construct for the heterologous expression of the decaheme MtrA from *S. oneidensis* MR-1 in *Synechocystis* sp. PCC6803 (hereafter *Synechocystis*). In addition to the growth, viability, and photosynthetic activity, the light-dependent exoelectrogenicity of the recombinant cells was also characterized using both colorimetric and electrochemical measurements. This demonstration not only provides a basis for genetically re-wiring a cell’s metabolism for photovoltaic applications but, more fundamentally, also illustrates a genetic approach to electronically couple the distinct light-harvesting metabolism of the living cells with external electrodes.

## Results and Discussion

### Expression of a MtrA-Flag fusion protein in *Synechocystis* sp. PCC6803

The expression of foreign c-type cytochromes in cyanobacteria poses significant challenges, as it requires post-translational modifications for covalent heme attachment as well as balanced expression levels that do not adversely affect the cell’s metabolism [30]. These challenges are exacerbated by the slower doubling rates and limited synthetic biology toolbox available for cyanobacteria compared to more genetically amenable microbes like *E. coli*, as well as the attempt to attach not one, but multiple hemes per protein, a feat that requires the allocation of additional metabolic resources. *Synechocystis* was ultimately chosen as the cyanobacterial host strain because of the relative ease with which the strain is able to uptake exogenous DNA and the relatively larger synthetic toolbox available for genetic engineering of this model organism.

To optimize *mtrA* transcription and translation, a set of new expression vectors was created that harbored a redesigned *mtrA* sequence adjusted for *Synechocystis* codon usage (IDT codon optimization tool). This sequence was fused to a flag-tag for sensitive analytical detection. Initial trials with inducible promoters did not lead to detectable expression of MtrA (data not shown). However, detectable expression levels were ultimately achieved using a strong cpc560 promoter [39]. Protein expression was compared using two different signaling sequences, the native *mtrA* signaling sequence from *S. oneidensis mtrA* and the native futA signaling sequence from *Synechocystis* [40]. This strategy led to the two expression constructs depicted in **Figure 2A**. The constructed plasmids were sequenced, and the confirmed sequences are provided in **Table S1**. The plasmids were individually transformed into wildtype (wt) *Synechocystis* through triparental mating conjugation, and the transformation was confirmed using colony PCR (**Figure S1A**).

**Figure 2.**
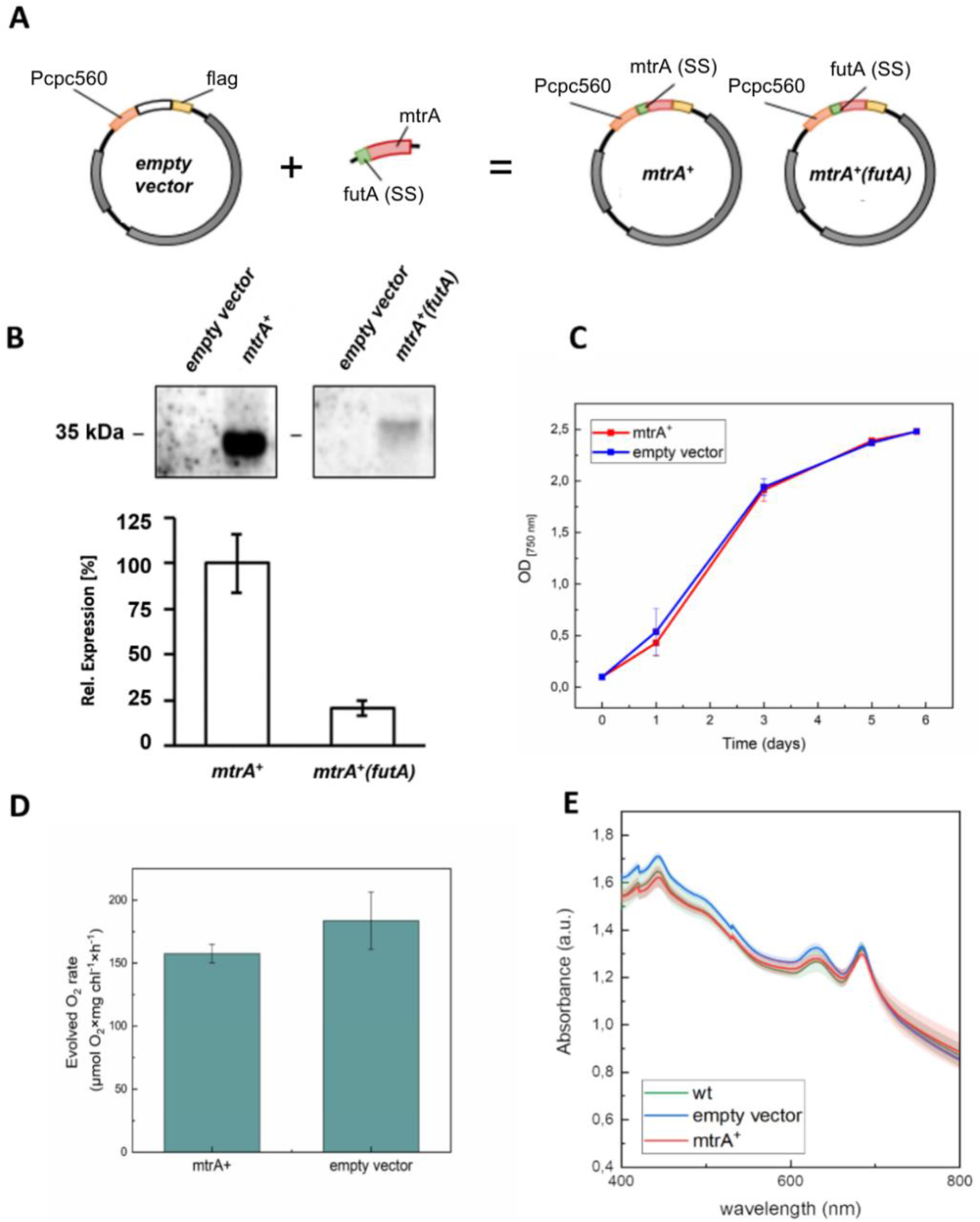
Expression of a MtrA-Flag fusion protein in *Synechocystis* sp. PCC6803. (**A**) Scheme of the expression vectors harboring distinct signaling sequences (SS). Corresponding sequences are provided in **Table S1**. (**B**) Top: Western Blot analysis of *Synechocystis* protein extracts using α-FLAG antibody and chemilumenescence substrates; Bottom: quantitative comparison of MtrA-Flag signals from constructs harboring either the *S. oneidensis* MtrA signaling sequence (mtrA^+^) or the *Synechocystis* futA signaling sequence (mtrA^+^(futA)). (**C**) Growth curves of three biological triplicates of mtrA+ (red) and empty vector control (blue) strains and their respective based on optical density measurements at 750 nm absorption (OD_750_). (**D**) Oxygen production rate of mtrA^+^ (red) and empty vector control cells (blue) under illuminated condition: 200 μmol photons m^-2^ s^-1^. (**E**) Absorption spectra of MtrA-expressing, wt, and empty vector control strains.

MtrA expression was confirmed by Western-blotting and the immunostaining of protein extracts (**Figure 2B** and **Figure S1B**). However, the construct with the futA signaling sequence led to a more than 4-fold reduction in MtrA-Flag (∼35 kDa) expression (bottom **Figure 2B**). Since futA is a highly abundant periplasmic protein, the low expression levels achieved using the corresponding signaling sequence were unexpected and are hypothesized to stem from an unfavorable sequence combination that adversely affects RNA stability or translation. Because of the lower expression level, only the strains encoding the intrinsic *mtrA* signaling sequence from *S. oneidensis* were used for further experiments. Transformed strains were grown under different light conditions to find favorable expression conditions, and fractional protein extracts were used to confirm to localization of the MtrA-Flag (**Figure S1C**). The fusion protein was neither detected in the soluble protein fraction nor could it be solubilized from the membrane fraction. These observations suggest that MtrA may behave as an insoluble protein when expressed in *Synechocystis* in absence of the MtrB, in agreement with recent observations that have shown MtrA localization within MtrB [12, 41].

The effects of MtrA expression on cell growth and photosynthetic activity was further studies through comparison of growth curves and oxygen evolution. It was observed that the heterologous expression of mtrA does not pose a severe metabolic burden for *Synechocystis*. As shown in **Figure 2C**, the MtrA-expressing cells (mtrA^+^) showed a nearly complete overlap of OD_750_ growth curves compared to empty vector control cells, indicating that the decaheme protein expressions at these levels did not significantly impair cell growth. However, a comparison of the oxygen evolutions indicated a slight decrease in oxygen evolution for the MtrA-expressing cells (157.58 ± 7.30 μmol O_2_×mg chl^-1^×h^-1^) compared to the empty vector control cells (183.64 ± 22.69 μmol O_2_×mg chl^-1^×h^-1^), though this difference was found to lie within measurement error (**Figure 2D**). In addition, a comparison of the absorption spectra of mtrA^+^, empty vector and wt strains under standard growth conditions indicates no significant difference in the photosynthetic pigment content (**Figure 2E**). In particular, all the curves show the expected presence of the three characteristic absorption peaks assigned to (i) the chlorophyll (at 436 nm), (ii) the photosynthetic systems (680 nm) and (iii) the phycocyanin (PC) pigments (at 625 nm) [42]. These results therefore indicate no significant difference in growth and photosynthetic activity of the growth of MtrA-expressing cells under the tested expression and growth conditions.

### Colorimetric characterization of the electrogenic activity of MtrA-expressing *Synechocystis*

Cyanobacteria are model organisms for studying photosynthesis and related applications such as biofuel synthesis and electricity generation [43-48, 1, 3-6]. In these organisms, photosynthesis takes place in the thylakoid membranes (TMs), which contain all the complexes involved in the photosynthetic electron transfer chain (PETC) (i.e., the photosystem I and II [PSI, PSII] with their reaction centers, and cytochrome *b*_*6*_*f*) [49] (**Figure 3A**). Upon illumination, PSII uses solar energy to split water and to reduce the plastoquinone (PQ) pool. The extracted electrons are then transported from the PQ pool to the cytochrome *b*_*6*_*f* complex before being transferred to a soluble electron carrier on the luminal side of the thylakoid membrane. This carrier reduces the PSI reaction center (chlorophyll [P700]), which becomes re-oxidized by ferredoxin-NADP reductase to produce nicotinamide adenine dinucleotide phosphate (NADPH). The photogenerated electrons are ultimately transferred to the periplasm through a putative trans-membrane protein (TMP) on re-oxidation of the NADPH. [49-50]

**Figure 3.**
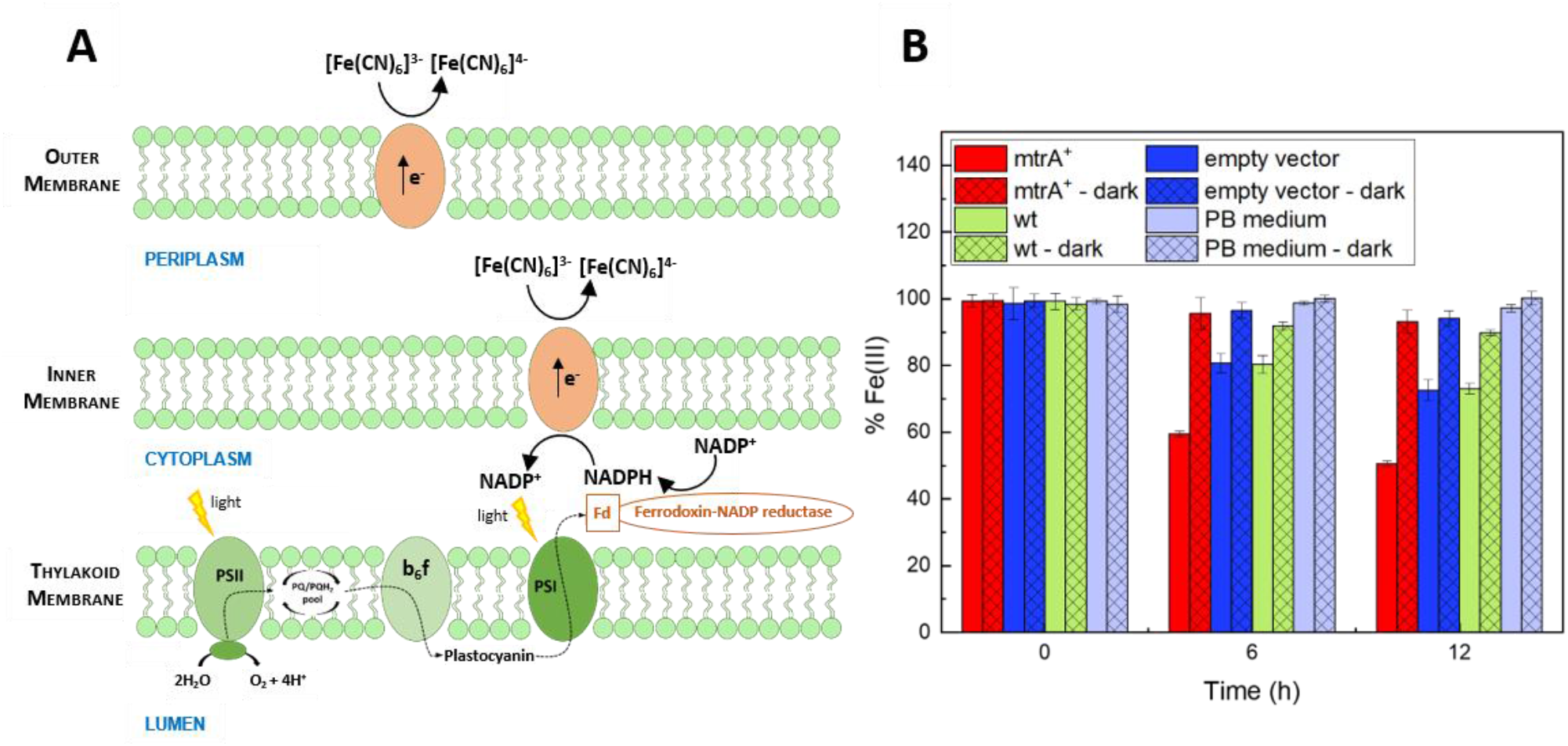
Periplasmic electrogenic activity of MtrA-expressing *Synechocystis*. (**A**) Scheme of PETC in cyanobacteria. The schematic depicts the interaction of ferricyanide [Fe(CN)_6_]^3-^ with outer membrane proteins as well as inner membrane proteins in the periplasmic region. (**B**) Colorimetric assay for monitoring cell electrogenicity over time. The assay was applied to mtrA^+^ cells (red), cells incubated with empty vector (blue), and wt control cells (green). Periplasmic electrogenic activity was quantified by spectrophotometrically measuring Fe(III) content (in percentage %) at 420 nm under dark (with shading) and light (without shading) conditions. Lilac bars represent Fe(III) content in a phosphate buffer (PB) control solution containing 1 mM K_3_Fe(CN)_6_.

Ferricyanide [Fe(CN)_6_]^3-^ is known to penetrate the outer membrane of cyanobacteria, as schematically depicted in **Figure 3A**. Consequently, [Fe(CN)_6_]^3-^ is often used in biological photovoltaics to extract photogenerated electrons from the microbe’s periplasm [51]. On reduction, [Fe(CN)_6_]^3-^ is converted to ferrocyanide [Fe(CN)_6_]^4-^, resulting in a decrease in absorption to 420 nm [52]. The periplasmic electrogenic activities of the wt *Synechocystis*, MtrA^+^, and empty vector control strains can therefore be compared by monitoring the change in absorption at 420 nm of cell filtrates in the presence of [Fe(CN)_6_]^3-^. Based on this assay, all tested strains showed increased electrogenicity over time under illuminated conditions compared to dark conditions in the presence of 1 mM [Fe(CN)_6_]^3-^ (**Figure 3B**). Under dark conditions, all the strains showed similar changes in absorption over time. These observations are in agreement with previous studies that have reported enhanced electrogencity from photosynthetically derived electrons under light conditions compared to the residual electrogenicity observed under dark conditions that is largely attributed to respiration [53]. Interestingly, and in contrast to the dark conditions, the tested strains showed variable electrogencity when under illumination. In particular, the MtrA-expressing cells showed a ca. 50% reduction of the initial Fe(III) concentration compared to the ca. 20% observed for the wt and empty vector controls strains respectively after 12 h. By comparison, no change in absorption was observed for media in the absence of cells, neither under dark nor light conditions. Considering the comparable photosynthetic activities measured for the different strains (see **Figure 2D**), these results indicate increased accessibility of photosynthetically derived electrons in the periplasm and/or outer membrane of the MtrA-expressing strain.

### Electrochemical characterization of electrogenic activity of MtrA-expressing *Synechocystis*

The light responses of MtrA-expressing *Synechocystis*, wt, and empty vector control cells were investigated by performing chronoamperometric (CA) and cyclic voltammetry (CV) measurements in a single chamber, three-electrode electrochemical cell. The electrochemical cell, schematically shown in **Figure 4A**, consisted of a working electrode (WE), a counter electrode (CE) and a reference electrode (RE). The WE was composed of a graphite-based membrane electrode [54-56] (**Figure 4A**, left). The graphite rode was cleaned in an ultrasonic bath prior to addition of the cells, and a dialysis membrane was added to confine the cells in close proximity to the electrode and to prevent their detachment. A Pt wire and standard Ag/AgCl (vs. KCl) electrode were employed as counter and reference electrodes, respectively. The electrolyte consisted of a freshly prepared phosphate buffer solution supplemented with 10 mM NaCl and 5 mM MgCl_2_ (PB) containing 1 mM of K_3_Fe(CN)_6_, the latter of which was used to shuttle electrons from the periplasm and outer membrane to the electrode.

**Figure 4.**
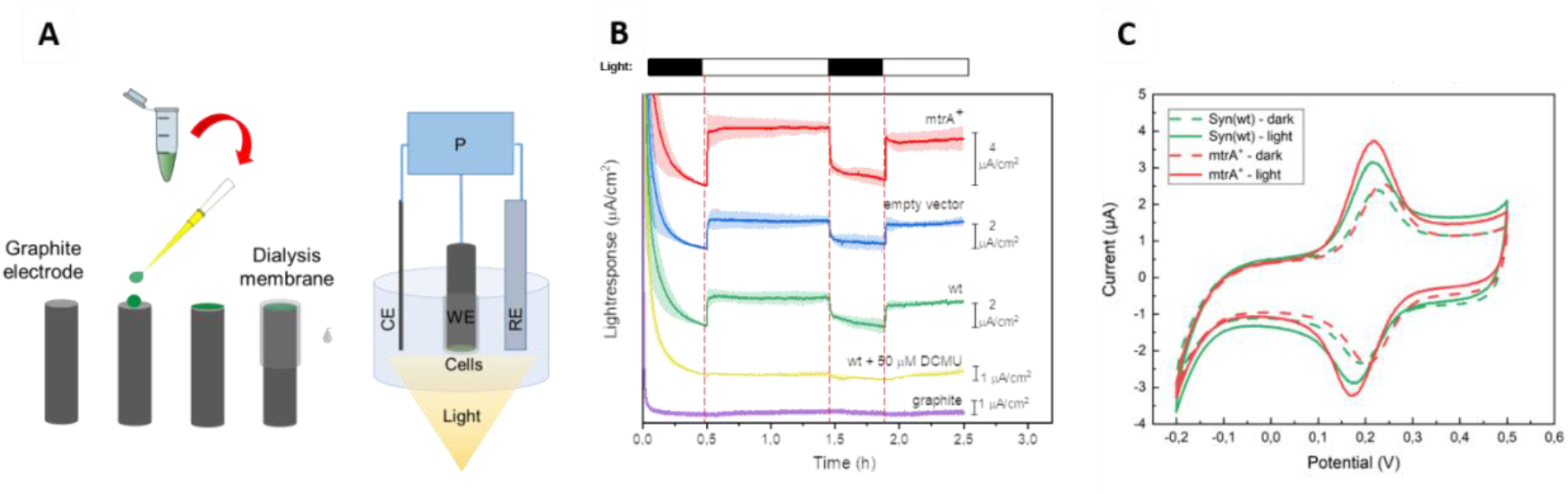
Electrochemical characterization of MtrA-expressing *Synechocystis*. (**A**) Schematization of the *Synechocystis*-modified graphite membrane electrodes (left) and of the electrochemical set-up (right). (**B**) CA measurements of mtrA^+^ cells (red), empty vector cells (blue), wt cells (green), wt cells incubated for 15 min in DCMU (yellow), and unmodified graphite electrode (purple) in the presence of 1 mM Fe(CN)_6_ during multiple light-dark cycles. Measurements were taken at an applied potential of 300 mV. (**C**) Representative CVs of wt- and mtrA^+^- expressing cells (green and red, respectively) immobilized on graphite working electrode in the presence of 1 mM K_3_Fe(CN)_6_ under dark (dotted lines) and light (bold lines) conditions. Measurements were taken using a sweep rate of 5 mV/s. All the CA measurements were taken using at least three biological replicates. The bold line corresponds to the mean value of the current density, and the shaded regions represent the standard deviation (SD).

The potentiostatic CA measurements were taken during dark-light cycles at an applied oxidative potential of 300 mV, corresponding to the oxidation potential of the added [Fe(CN)_6_]^4-^ as measured by CV (**Figure S2A**). For these CA measurements, the cells were incubated in the dark for 30 minutes, followed by 1 hour incubation under 500 μmol m^-2^s^-1^ of LED light illumination, 30 minutes of dark incubation, and another 30 minutes under the light conditions (**Figure 4B**, individual CA measurements are shown in **Figure S2 B-D**). During the first 30 minutes, all cells showed an initial decrease in current, indicative of the diffusion-limited extraction of capacitive charge reported in previous studies [57]. When illuminated, the cells showed an expected increase in the current density, reaching a plateau after approximately 5 minutes. The empty vector and wt control cells showed a comparable photoresponse, achieving stabilized current densities of about (2.1 ± 0.5) μA/cm^2^ and (2.1 ± 0.3) μA/cm^2^, respectively, after one hour of light exposure. The observed photoresponses were reversible, diminishing to baseline levels when the light was turned off and recovering on illumination. These observations are in agreement with previous ferricyanide-based electrochemical characterizations of wt *Synechocystis* cells [48]. In comparison to both the wt and empty vector cells, the MtrA-expressing *Synechocystis* cells showed a two-fold increase in current density, achieving a stabilized photocurrent of (4.3 ± 1.6) μA/cm^2^, in agreement with the enhanced electrogenicity observed with the colorimetric ferricyanide assay presented in **Figure 3B**. By contrast, no photocurrent was observed for control measurements in the absence of cells, attributing the observed response to the light-induced activity of the cells rather than a Fe(CN)_6_^3^ photoresponse. To examine the contributions of photosynthetic activity to the photocurrent, additional control measurements were performed in the presence of 3-(3,4-dichlorophenyl)-1,1-dimethylurea (DCMU), a renown photosynthesis inhibitor [51]. After 15 minutes of incubation in 50 μM DCMU, the wt cells showed an 80% reduction in photocurrent, similar to the percentage of reduction in photosynthetic activity reported in previous studies [49]. These results suggest that the improved electrogenicity of the MtrA-expressing strain can mostly be attributed to the enhanced extraction of electrons derived from photosynthesis.

In addition to the CA characterizations, CV measurements of the wt and MtrA-expressing cells were taken after 30 minutes of incubation in the dark and after 30 minutes of 500 μmol m^-2^s^-1^ illumination (**Figure 4C**). The measurements were taken at a sweep rate of 5 mV/s, which previously has shown to be sufficient in kinetically capturing extracted electrons from the photosynthetic metabolism of the cell [57]. As shown in **Figure 4C**, in the presence of 1 mM K_3_Fe(CN)_6,_ both wt and mtrA^+^ cells show a notable increase in the anodic current when under illumination compared to dark conditions. In addition, all the curves show sufficient reversibility in terms of peak current ratios (i_pa_/i_pc_ ≈ 1). These observations are in agreement with previous reports [57]. However, whereas both wt and mtrA^+^ cells show comparable capacities under dark conditions, the mtrA^+^ cells exhibit a 30% increase in the anodic (oxidation) current peak compared to the wt cells. These measurements further support a mechanism based on the enhanced extraction of photosynthetically derived electrons for the MtrA-expressing cells.

The enhanced photocurrent of the mtrA^+^ cells compared to wt cells is attributed to MtrA expression, which is expected to facilitate indirect-as opposed to direct-EET (see **Figure 4A**). The latter mechanism based on direct EET would otherwise require the co-expression of outer membrane proteins, such as MtrB and MtrC, or OmcS, as demonstrated in a previous study [29], to enable directed electron transfer from the outer membrane to the electrode surface. In agreement with the proposed mechanism of EET, the wt and MtrA-expressing cells show only a slight increase in current on illumination in the absence of mediator (**Figure S3**). This diminished photoresponse is attributed to the insulating outer membrane. In addition, under mediatorless conditions, both the wt and MtrA-expressing cells exhibit similar currents (**Figure S3**), indicating that the enhancement in photocurrent for the MtrA cells under mediated conditions occurs largely from inside the cell, and not from the outer membrane where the MtrA is not expected to localize.

The stability of the system was investigated by performing long-term CA experiments using 1 mM K_3_Fe(CN)_6_ as an electron mediator. Subsequent 1 hour light–dark cycles were conducted and the CAs lasted up to 24 hours in a single chamber of electrochemical setup (**Figure S3**) and up to 13 hours in a H-shaped double chamber device (**Figure 5A**). For the double chambered setup a carbon felt cathode electrode immersed in a PB buffer supplemented with 10 mM K_3_Fe(CN)_6_ has been employed. In both cases, although over multiple light cycles a decreased of the achieved current densities was observed, mtrA^+^ cells showed retained improved EET capability as compared to empty vector control cells. This decay was previously reported (55-58) when an electrolyte lacking nutrients and external carbon sources was used, and it might also be associated with the oxygen production at the anode resulting in current losses in absence of N_2_ purging (37).

**Figure 5.**
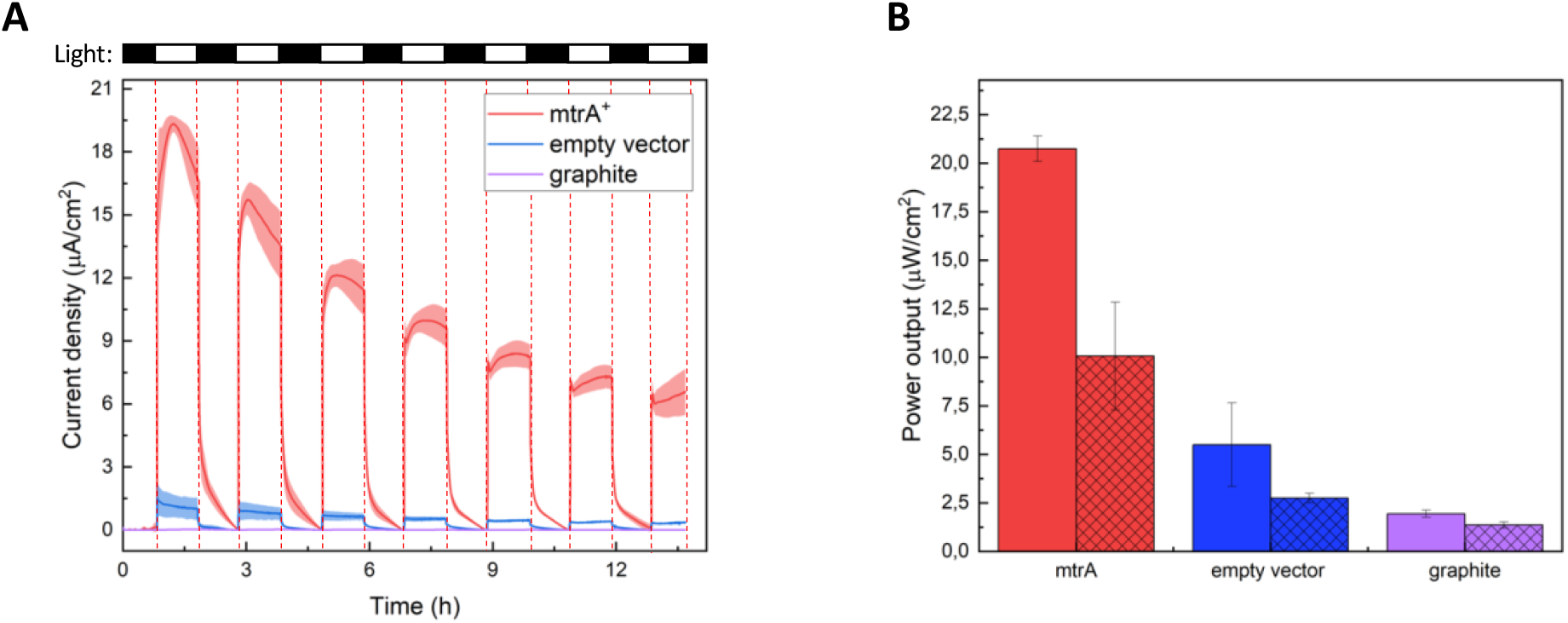
Implementation of MtrA-expressing *Synechocystis* in a BPV device. (**A**) Long-term CA measurements of mtrA^+^ cells (red), empty vector cells (blue) and unmodified graphite electrodes (purple) in the presence of 1 mM K_3_Fe(CN)_6_ during multiple light-dark cycles. Measurements were taken at an applied potential of 300 mV using a BPV device (H-shaped double chamber electrochemical cell) equipped with a carbon felt as cathode, immersed in a PB electrolyte supplemented with 10 mM K_3_Fe(CN)_6_. (**B**) Peak power outputs produced by mtrA^+^ cells (red) and empty vector cells (blue) under dark (with shading) and light (without shading) conditions. Lilac bars represent the power output recorded using unmodified graphite anodes in a phosphate buffer (PB) control solution containing 1 mM K_3_Fe(CN)_6_.

Linear sweep voltammetry (LSV) measurements were conducted in the double chamber devices to estimate the peak power outputs achieved using the mutant strains. The corresponding derived power outputs expressed relative to the anode area are reported in **Figure 5B**. As expected, in light conditions MtrA-expressing cells showed a 4-fold improved power output compared to empty vector control cells. Interestingly, this latter showed power densities comparable to the ones previously reported by Bradley et al. (37) obtained using wild-type *Synechocystis*, confirming the similar extent of electrogenicity of wt and empty vector control cells we determined using the colorimetric assay (**Figure 3**). As BPV performances are influenced by several parameters (such as surface-to-volume ratio, electrodes materials, photosynthetic organism employed, geometry and internal resistance of the device, illumination sources and intensities), quantitative comparisons with the power outputs already reported in literature are hard to achieve. Nonetheless, it is worthwhile noting that the increase in peak power outputs produced by mtrA^+^ mutant cells in our system is comparable to the improvements obtained by engineered *Synechocystis* lacking three terminal oxidases (37), with the advantage of a retained higher viability.

## Conclusions

Heme proteins play a key role in regulating the redox metabolism of a cell. The introduction foreign cytochromes inside a cell can thus rewire the metabolism by re-directing electron flux. In this study, the heterologous expression of the decaheme MtrA was used to increase cell exoelectrogenicity through enhanced EET. This was achieved within a photosynthetic host to enable light-activated charge extraction from a living photovoltaic. Despite the low expression levels of this protein, as indicated by the relatively long exposure times required for the Western Blot imaging (150 s), the bioengineered cells show a measurable increase in exoelectrogenicity with retained photosythetic activity and cell viability. Future approaches to further increasing protein expression could include the exploration of alternative promoters with more controlled protein expression. In addition, protein expression could also possibly be improved by facilitating heme insertion through the concomitant overexpression of the cytochrome c machinery (ccm) in the presence of added heme precursors such as δ-aminolevulinic acid (δ-ALA). Such approaches for facilitating protein expression could be critical for reconstituting the entire MtrCAB pathway, which requires the co-expression of MtrB and MtrC. Based on previous studies in *E. coli* [28], the full reconstitution of this defined pathway from the natural exoelectrogen, *S. oneidensis*, is expected to increase indirect and direct EET.

Although this study focuses on charge extraction mechanisms in the framework of photovoltaics, previous studies have also utilized the MtrABC pathway in the reverse direction for microbial electrosynthesis [59, 60]. In these applications, the inward flux of electrons is used to drive the synthesis of high-value chemicals [61], including alcohols [59] and organic acids [62], by exploiting the complex networks of reactions within a cell metabolism. These advantages are exasperated with photosynthetic microbes, which are capable of harnessing solar energy to drive thermodynamically unfavorable reactions such as water-splitting. This ability for foreign cytochrome expression inside new hosts that benefit from such endogenous light-harvesting machinery therefore brings new possibilities for reaching efficient biological energy and sustainability technologies.

## Methods

### Cloning & plasmid construction

A cyanobacterial expression vector was designed with the strong *cpc560* promoter [39] and a 3xFLAG-tag for immunodetection. The empty vector was constructed by Gibson assembly with all fragments amplified using Phusion DNA polymerase (New England BioLabs). PCR products were extracted from agarose gels after electrophoresis and purified using the QIAEX II Gel Extraction Kits (Qiagen) according to the manufacturer’s instructions. The plasmid backbone was based on a pUR-vector [63] with a single SapI restriction site removed by site-directed mutagenesis using primers 1 and 2, as described elsewhere [64]. Primers 3-6 were used to amplify the plasmid as two separate parts in order to reduce fragment size. The *cpc560* promoter was amplified from *Synechocystis* genomic DNA (primers 7,8) and a SapI cloning site harboring a 430 nt stuffer sequence was amplified from plasmid pBSYA1 (primers 9,10). The purified PCR fragments were concentrated by ethanol precipitation and assembled according to the one-step isothermal DNA assembly protocol [65]. The reaction mixture was transformed into *E. coli* DH5α cells by electroporation and colonies were selected on LB agar containing 50 μg/ml kanamycin. Clones were screened by colony PCR, and the plasmid containing the empty vector (pNS02-empty) was isolated from positive clones. The correct assembly of the expression cassette was verified by sequencing (primers 11,12). Two overlapping DNA strands encoding a codon-optimized *mtrA* gene from *Shewanella* were chemically synthesized (Eurofins Genomics) and fused by overlap-extension PCR using flaking primers 13 and 14 containing a SapI restriction sites. The coding sequence was subcloned, excised by SapI digestion, purified by gel electrophoresis, and ligated into the SapI digested pNS02-empty plasmid. After transformation into *E. coli* DH5α, the plasmid containing the mtrA-expressing vector (pNS02-*mtrA)* was isolated from positive clones and verified by sequencing. The pNS02 vectors were transferred to *Synechocystis* cells via tri-parental mating using the *E. coli* strain ED8654 harboring the conjugative plasmid pRL443 as a helper strain.

**Table 1:**
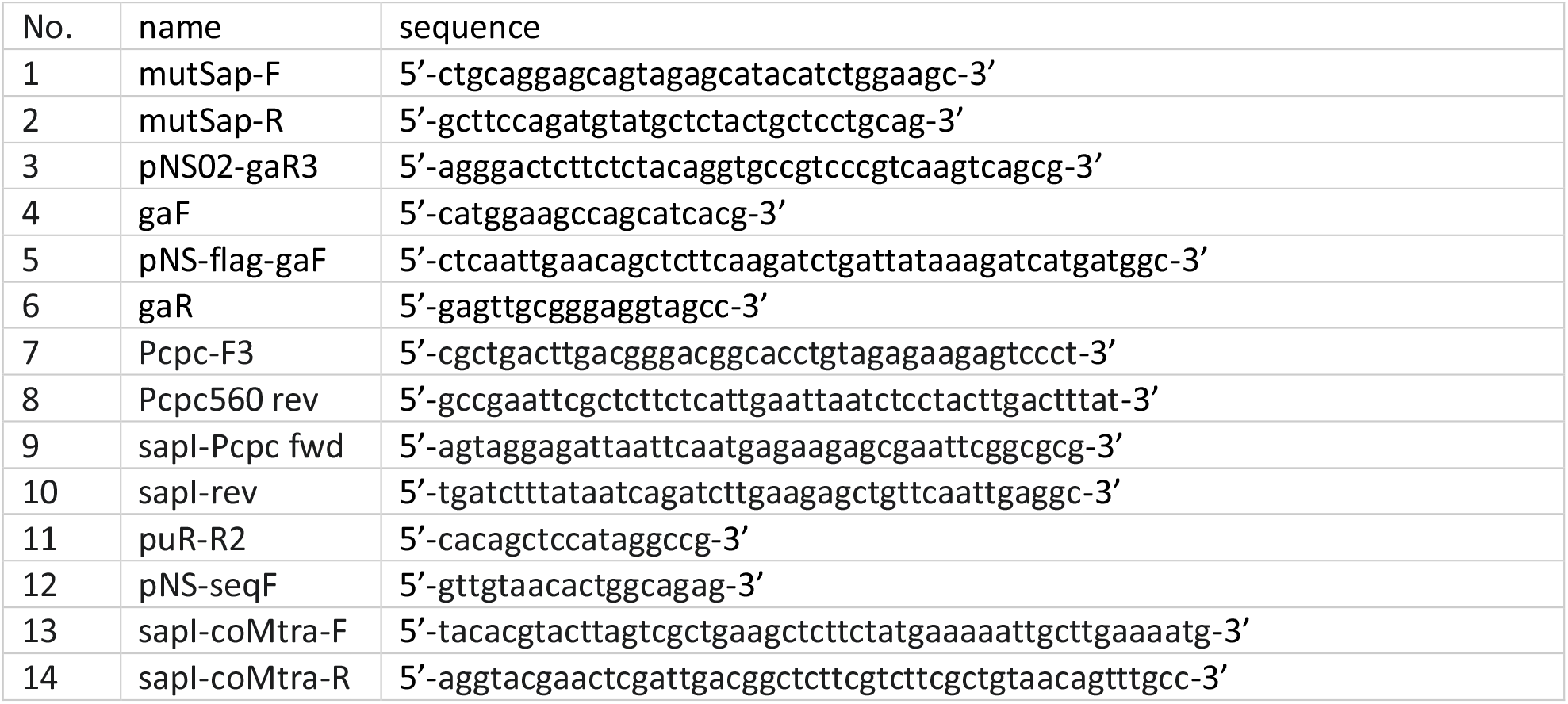
DNA oligonucleotides used in this study.

### Strains and growth conditions

We used a *Synechocystis* sp. PCC 6803 wild type sub-strain (PCC-M) that had been re-sequenced 2012 from the lab of S. Shestakov (Moscow State Forest University) [66]. Cultures were propagated on 0.75% agar plates containing BG11 medium [67] supplemented with 0.3% sodium thiosulfate, and liquid cultures were grown in BG11 supplemented with 10 mM Tris(hydroxymethyl)methyl-2-aminoethane-sulfonic acid (TES) buffer (pH 8.0) at 30°C under white LED light of 50 μmol photons m^−2^ s^−1^ and constant shaking (Infors HT Multitron Pro Shaker). Strains harboring pNS02 plasmids were supplemented with 50 μg/ml kanamycin.

### Protein expression and immunodetection

Liquid cultures of *Synechocystis* MtrA^+^ and control strains were diluted to an OD_750nm_ of 0.5 in 50 ml BG11 medium and grown for 24 h under standard conditions as described above. Cells were harvested by centrifugation (2.200 x g; 10 min; 4°C) and resuspended in 500 μl extraction buffer (50 mM HEPES (pH 7), 5 mM MgCl_2_, 25 mM CaCl_2_, 150 mM NaCl, 10% (v/v) Glycerol, 0.1% (w/v) Tween20) supplemented with 12.5 μl Protease Inhibitor Cocktail Bacteria (Carl Roth). After the addition of 500 μl acid-washed sand (0.1-0.3 mm), cells were lysed at 4°C by mixing in an LLG uniTEXER for 5 min at maximum speed. Sand and unbroken cells were removed by centrifuging two times (1500 x g; 2 min; 4°C), and the raw extract was separated in soluble and membrane/insoluble protein fractions by centrifugation (21.000 x g; 30 min; 4°C). Protein samples were diluted with SDS loading buffer, heated to 95°C for 5 min, briefly centrifuged, separated by 12% Laemmli SDS-PAGE (BioRAD MiniProtean), and transferred to PVDF membranes using Towbin buffer (25 mM Tris, 192 mM glycine, 20% (v/v) methanol, 0.1% SDS). Membranes were blocked with milk-powder (10% in TBS-T) and stained overnight at 4°C with Pierce FLAG-HRP conjugate 4°C (1:3.000 in TBS-T). MtrA-FLAG fusion proteins were visualized with Pierce Super Signal Femto ECL HRP substrate using a Fusion Imager (FUSION solo S, Vilber) with an exposure time of 150s.

### Ability of Fe(III) reduction

*Synechocystis* mtrA^+^, empty vector, and wt control cells were washed (centrifugation at 3200 x g for 10 min) and resuspended in 20 mL of phosphate buffer solution supplemented with 10 mM NaCl and 5 mM MgCl_2_ (PB) containing 1mM K_3_Fe(CN)_6_ to an OD_750nm_ of 2. Flasks were incubated at 30 °C under illumination (50 μmol photon m^-2^ s^-1^) for 12 hours. At regular 6-h intervals, 200-μL cell aliquots were filtered to measure the Fe(III) absorbance at 420nm using a Varioskan™ LUX multimode microplate reader (Thermo Scientific™). At each time point, the cell’s OD was simultaneously measured. Three biological triplicates were taken for each strain, and three distinctive measurements were taken for each triplicate at each time point (technical triplicates). The mean values shown herein were normalized by the measured OD and reported as mean values with their corresponding standard deviation. As an additional control, Fe(III) absorption and OD_750 nm_ measurements were also taken at each time point for flasks containing the assay medium in the absence of cells.

### Light response of MtrA-expressing *Synechocystis* on graphite electrodes

The light response of *Synechocystis* mtrA^+^, wt, and empty vector control cells were compared using CA and CV measurements in a single chamber, three-electrode electrochemical cell. Cells were immobilized on graphite WEs that were previously cleaned in an ultrasound bath (10 min ethanol + 10 min deionized H_2_O) and sterilized in an autoclave. A Pt wire was used as the CE and Ag/AgCl (vs. KCl) as the RE.

For the preparation of the WEs, cell suspensions (OD_750nm_ of 2 in 1 mL) were pelleted and washed in PB. After centrifugation at 5000 rpm for 5 min., the washed pellets were resuspended in 10 μL of PB buffer, drop-casted on the surface of a graphite rod electrode (AlfaAesar), and dried for 30 min. The resulting cell-modified graphite electrodes were encased in a dialysis membrane (14 kDa molecular weight pore size, Sigma-Aldrich) that was previously soaked in deionized water for 1 h before insertion in the electrochemical cell. The electrochemical cell was filled with PB containing 1 mM of K_3_Fe(CN)_6_. During the CA measurements, the system was left to stabilize in the dark for 30 min, followed by 1 h incubation under 500 μmol m^-2^s^-1^ of LED illumination, followed by another 30 min in the dark and another 30 min under the same light conditions. Control CA measurements were taken for bare graphite electrodes encased in the same dialysis membrane in the absence of cells, as well as wt-modified and mtrA^+^-modified electrodes soaked for 15 min in 50 μM DCMU (Sigma-Aldrich). At least three different biological triplicates were taken for each sample, and the results shown herein represent the mean values with the corresponding standard deviations. For the CV measurements, measurements were taken at a scan rate of 5 mV/s under dark conditions 30 min after the construction of the electrochemical cell in dark. Upon illumination, the system was left to stabilize for 30 min, and the voltammograms were then recorded under 500 μmol m^-2^s^-1^ of LED illumination. Power outputs produced by MtrA-expressing cells and empty vector control cells were derived performing LSV measurements at a scan rate of 5 mV/s using a H-shaped double chamber electrochemical cells. The anode chambers were equipped with the graphite rod electrodes prepared as previously mentioned and an Ag/AgCl reference electrode immersed in 100 mL of PB electrolyte supplemented with 1 mM K_3_Fe(CN)_6_. Carbon felts of 0.25 cm^3^ were used as cathodes and immersed in PB catholyte containing 10 mL of 10 mM K_3_Fe(CN)_6_. The two chambers were separated by a Nafion membrane (Chemie Brunschwig AG). LSV in dark condition were performed at the end of CA experiments lasted for 14 hours after letting stabilize the system for 1 hour. Open circuit potential (OCP) measurements in light condition (∼500 μmol m^-2^s^-1^ intensity) were afterwards conducted for 1 hour prior to record LSV curve under illumination. The peak power outputs were then extracted from the LSV curves using the PSTrace 5.2 software (Palmsens).

## Acknowledgements

The authors are thankful for support from the Swiss National Science Foundation (project number PYAPP2_154269/1 and IZLIZ2_182972).

