## supplementary information for "Living Photovoltaics based on Recombinant Expression of MtrA Decaheme in Photosynthetic Bacteria"

[1] Institute of Chemical Sciences and Engineering (ISIC), École Polytechnique Fédérale de Lausanne (EPFL), 1015-Lausanne, Switzerland

### *Supplementary Information*

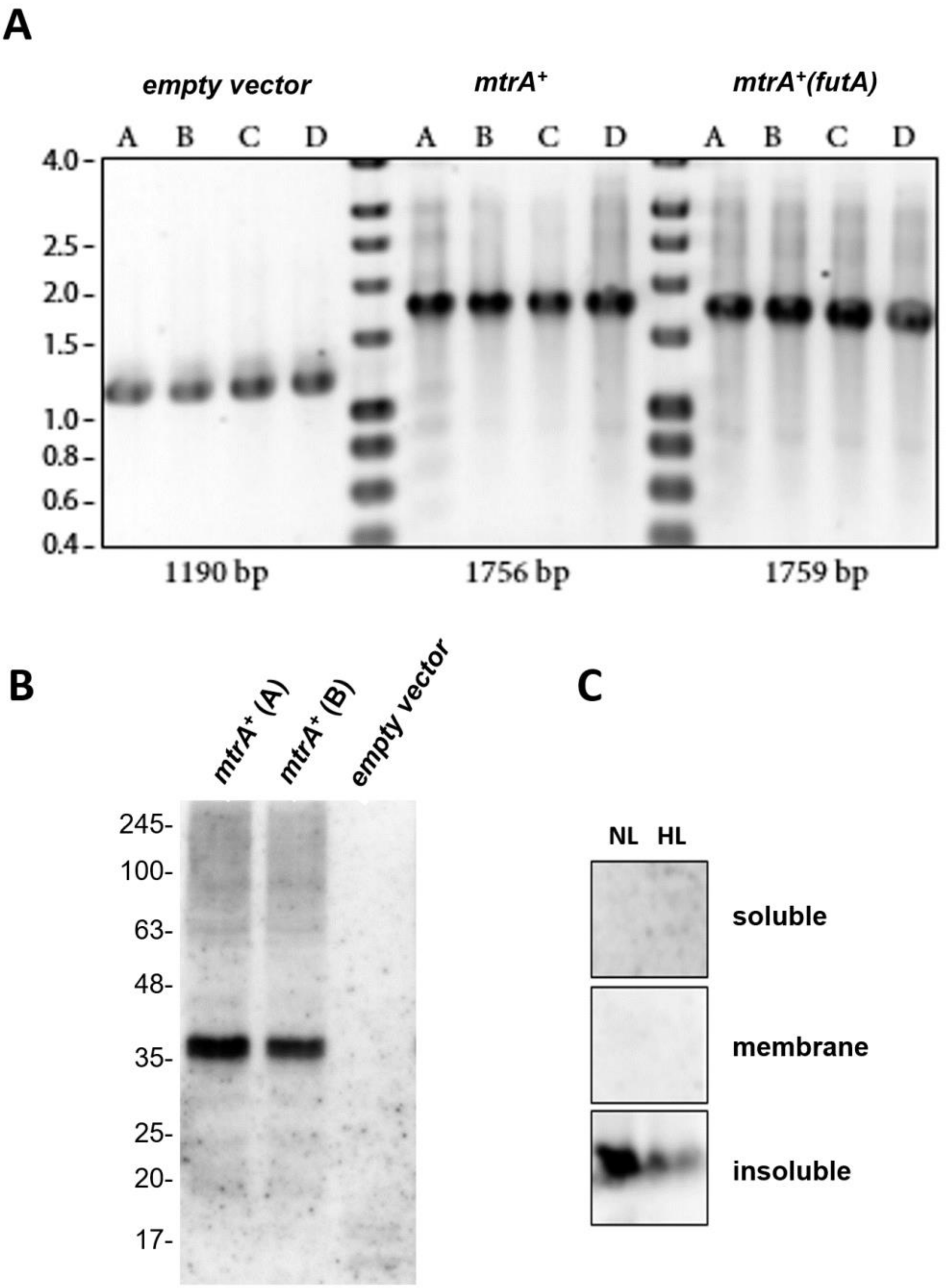


**Figure S1: (A)** Colony PCR and gel electrophoresis on 0.75% agarose gels used to verify the conjugative transfer of the expression plasmids or empty vectors into *Synechocystis* (four colonies each). **(B)** Western Blot of protein extracts from cell cultures grown under standard illumination (50 µmol photons $m^{-2}$ $s^{-1}$). Proteins were separated by 12% SDS-PAGE, transferred to PDVF membranes and stained with an α-FLAG antibody. **(C)** Western Blot analysis of different protein fractions from cells grown either under 50 (NL) or 200 (HL) µmol photons $m^{-2}s^{-1}$ light conditions.

**Table S1: Expression vector sequences**

| **MtrA$^+$** |
|---|
| agatctgattataaagatcatgatggcgattataaagatcatgatattgattataaagatgatgatgataaataataaggatccaacgctcggttgccgccg |
| ggcgtttttattctgcaggagcagtagagcatacatctggaagcaaagccaggaaagcggcctatggagctgtgcggcagcgctcagtaggcaatttttca |
| aaatattgttaagccttttctgagcatggtatttttcatggtattaccaattagcaggaaaataagccattgaatataaaagataaaaatgtcttgtttacaat |
| agagtgggggggtcagcctgccgccttgggccgggtgatgtcgtacttgcccgccgcgaactcggttaccgtccagcccagcgcgaccagctccggcaac |
| gcctcgcgcacccgctggcggcgcttgcgcatggtcgaaccactggcctctgacggccagacatagccgcacaaggtatctatggaagccttgccggttttg |
| ccggggtcgatccagccacacagccgctggtgcagcaggcgggcggtttcgctgtccagcgcccgcacctcgtccatgctgatgcgcacatgctggccgcc |
| acccatgacggcctgcgcgatcaaggggttcagggccacgtacaggcgcccgtccgcctcgtcgctggcgtactccgacagcagccgaaacccctgccgct |
| tgcggccattctgggcgatgatggataccttccaaaggcgctcgatgcagtcctgtatgtgcttgagcgccccaccactatcgacctctgccccgatttccttt |
| gccagcgcccgatagctacctttgaccacatggcattcagcggtgacggcctcccacttgggttccaggaacagccggagctgccgtccgccttcggtcttg |
| ggttccgggccaagcactaggccattaggcccagccatggccaccagcccttgcaggatgcgcagatcatcagcgcccagcggctccgggccgctgaactc |
| gatccgcttgccgtcgccgtagtcatacgtcacgtccagcttgctgcgcttgcgctcgccccgcttgagggcacggaacaggccgggggccagacagtgcgc |
| cgggtcgtgccggacgtggctgaggctgtgcttgttcttaggcttcaccacggggcaccccccttgctcttgcgctgcctctccagcacggcgggcttgagcac |
| cccgccgtcatgccgcctgaaccaccgatcagcgaacggtgcgccatagttggccttgctcacaccgaagcggacgaagaaccggcgctggtcgtcgtcca |
| caccccattcctcggcctcggcgctggtcatgctcgacaggtaggactgccagcggatgttatcgaccagtaccgagctgccccggctggcctgctgctggtc |
| gcctgcgcccatcatggccgcgcccttgctggcatggtgcaggaacacgatagagcacccggtatcggcggcgatggcctccatgcgaccgatgacctggg |
| ccatggggccgctggcgttttcttcctcgatgtggaaccggcgcagcgtgtccagcaccatcaggcggcggccctcggcggcgcgcttgaggccgtcgaacc |
| actccggggccatgatgttgggcaggctgccgatcagcggctggatcagcaggccgtcagccacggcttgccgttcctcggcgctgaggtgcgccccaagg |
| gcgtgcaggcggtgatgaatggcggtgggcgggtcttcggcgggcaggtagatcaccgggccggtgggcagttcgcccacctccagcagatccggcccgc |
| ctgcaatctgtgcggccagttgcagggccagcatggatttaccggcaccaccgggcgacaccagcgccccgaccgtaccggccaccatgttgggcaaaac |
| gtagtccagcggtggcggcgctgctgcgaacgcctccagaatattgataggcttatgggtagccattgattgcctcctttgcaggcagttggtggttaggcgc |
| tggcggggtcactaccccgccctgcgccgctctgagttcttccaggcactcgcgcagcgcctcgtattcgtcgtcggtcagccagaacttgcgctgacgcat |
| ccctttggccttcatgcgctcggcatatcgcgcttggcgtacagcgtcagggctggccagcaggtcgccggtctgcttgtccttttggtctttcatatcagtcac |
| cgagaaacttgccggggccgaaaggcttgtcttcgcggaacaaggacaaggtgcagccgtcaaggttaaggctggccatatcagcgactgaaaagcggc |
| cagcctcggccttgtttgacgtataaccaaagccaccgggcaaccaatagcccttgtcacttttgatcaggtagaccgaccctgaagcgctttttttcgtattcc |
| ataaaaccccccttctgtgcgtgagtactcatagtataacaggcgtgagtaccaacgcaagcactacatgctgaaatctggcccgcccctgtccatgcctcgct |
| ggcggggtgccggtgcccgtgccagctcggcccgcgcaagctggacgctgggcagacccatgaccttgctgacggtgcgctcgatgtaatccgcttcgtgg |
| ccgggcttgcgctctgccagcgctgggctggcctcggccatggccttgccgatttcctcggcactgcggccccggctggccagcttctgcgcggcgataaagt |
| cgcacttgctgaggtcatcaccgaagcgcttgaccagcccggccatctcgctgcggtactcgtccagcgccgtgcgccggtggcggctaagctgccgctcgg |
| gcagttcgaggctggccagcctgcgggccttctcctgctgccgctgggcctgctcgatctgctggccagcctgctgcaccagcgccgggccagcggtggcgg |
| tcttgcccttggattcacgcagcagcacccacggctgataaccggcgcgggtggtgtgcttgtccttgcggttggtgaagcccgccaagcggccatagtggc |
| ggctgtcggcgctggccgggtcggcgtcgtactcgctggccagcgtccgggcaatctgcccccgaagttcaccgcctgcggcgtcggccaccttgacccatg |
| cctgatagttcttcgggctggtttccactaccagggcaggctcccggccctcggctttcatgtcatccaggtcaaactcgctgaggtcgtccaccagcaccag |
| accatgccgctcctgctcggcgggcctgatatacacgtcattgccctgggcattcatccgcttgagccatggcgtgttctggagcacttcggcggctgaccatt |
| cccggttcatcatctggccggtggtggcgtccctgacgccgatatcgaagcgctcacagcccatggccttgagctgtcggcctatggcctgcaaagtcctgtc |
| gttcttcatcgggccaccaagcgcagccagatcgagccgtcctcggttgtcagtggcgtcaggtcgagcaagagcaacgatgcgatcagcagcaccaccgt |
| aggcatcatggaagccagcatcacggttagccatagcttccagtgccacccccgcgacgcgctccgggcgctctgcgcggcgctgctcacctcggcggcta |
| cctcccgcaactctttggccagctccacccatgccgcccctgtctggcgctgggctttcagccactccgccgcctgcgcctcgctggcctgctgggtctggctc |
| atgacctgccgggcttcgtcggccagtgtcgccatgctctgggccagcggttcgatctgctccgctaactcgttgatgcctctggatttcttcactctgtcgatt |
| gcgttcatggtctattgcctcccggtattcctgtaagtcgatgatctgggcgttggcggtgtcgatgttcagggccacgtctgcccggtcggtgcggatgcccc |
| ggccttccatctccaccacgttcggccccaggtgaacaccgggcaggcgctcgatgccctgcgcctcaagtgttctgtggtcaatgcgggcgtcgtggccag |
| cccgctctaatgcccggttggcatggtcggcccatgcctcgcgggtctgctcaagccatgccttgggcttgagcgcttcggtcttctgtgccccgcccttctccg |
| gggtcttgccgttgtaccgcttgaaccactgagcggcgggccgctcgatgccgtcattgatccgctcggagatcatcaggtggcagtgcgggttctcgccgcc |
| accggcatggatggccagcgtatacggcaggcgctcggcaccggtcaggtgctgggcgaactcggacgccagcgccttctgctggtcgagggtcagctcg |
| accggcagggcaaattcgacctccttgaacagccgcccattggcgcgttcatacaggtcggcagcatcccagtagtcggcgggccgctcgacgaactccgg |
| catgtgcccggattcggcgtgcaagacttcatccatgtcgcgggcatacttgccttcgcgctggatgtagtcggccttggccctggccgattggccgcccgac |
| ctgctgccggttttcgccgtaaggtgataaatcgccatgctgcctcgctgttgcttttgcttttcggctccatgcaatggccctcggagagcgcaccgcccgaa |
| gggtggccgttaggccagtttctcgaagagaaaccggtaagtgcgccctcccctacaaagtagggtcgggattgccgccgctgtgcctccatgatagcctac |
| gagacagcacattaacaatggggtgtcaagatggttaaggggagcaacaaggcggcggatcggctggccaagctcgaagaacaacgagcgcgaatcaa |
| tgccgaaattcagcgggtgcgggcaagggaacagcagcaagagcgcaagaacgaaacaaggcgcaaggtgctggtgggggccatgattttggccaagg |
| tgaacagcagcgagtggccggaggatcggctcatggcggcaatggatgcgtaccttgaacgcgaccacgaccgcgccttgttcggtctgccgccacgccag |
| aaggatgagccgggctgaatgatcgaccgagacaggccctgcggggctgcacacgcgcccccacccttcgggtagggggaaaggccgctaaagcggcta |
| aaagcgctccagcgtatttctgcggggtttggtgtggggtttagcgggctttgcccgcctttccccctgccgcgcagcggtggggcggtgtgtagcctagcgc |
| agcgaatagaccagctatccggcctctggccgggcatattgggcaagggcagcagcgccccacaagggcgctgataaccgcgcctagtggattattcttag |
| ataatcatggatggatttttccaacaccccgccagcccccgcccctgctgggtttgcaggtttgggggcgtgacagttattgcaggggttcgtgacagttattg |

caggggggcgtgacagttattgcaggggttcgtgacagttagtacgggagtgacgggcactggctggcaatgtctagcaacggcaggcatttcggctgagg
gtaaaagaactttccgctaagcgatagactgtatgtaaacacagtattgcaaggacgcggaacatgcctcatgtggcggccaggacggccagccgggatc
gggatactggtcgttaccagagccaccgacccgagcaaacccttctctatcagatcgttgacgagtattacccggcattcgctgcgcttatggcagagcagg
gaaaggaattgccgggctatgtgcaacgggaatttgaagaatttctccaatgcgggcggctggagcatggctttctacgggttcgctgcgagtcttgccacg
ccgagcacctggtcgctttcagctgtaatccgggcagcgcaacggaacattcatcagtgtaaaaatggaatcaataaagccctgcgcagcgcgcagggtca
gcctgaatacgcgtttaatgaccagcacagtcgtgatggcaaggtcagaatagcgctgaggtctgcctcgtgaagaaggtgttgctgactcataccaggcct
gaatcgccccatcatccagccagaaagtgagggagccacggttgatgagagctttgttgtaggtggaccagttggtgattttgaacttttgctttgccacgga
acggtctgcgttgtcgggaagatgcgtgatctgatccttcaactcagcaaaagttcgatttattcaacaaagccacgttgtgtctcaaaatctctgatgttaca
ttgcacaagataaaaatatatcatcatgaacaataaaactgtctgcttacataaacagtaatacaaggggtgttatgagccatattcaacgggaaacgtctt
gctcgaggccgcgattaaattccaacatggatgctgatttatatgggtataaatgggctcgcgataatgtcgggcaatcaggtgcgacaatctatcgattgt
atgggaagcccgatgcgccagagttgtttctgaaacatggcaaaggtagcgttgccaatgatgttacagatgagatggtcagactaaactggctgacggaa
tttatgcctcttccgaccatcaagcattttatccgtactcctgatgatgcatggttactcaccactgcgatccccgggaaaacagcattccaggtattagaaga
atatcctgattcaggtgaaaatattgttgatgcgctggcagtgttcctgcgccggttgcattcgattcctgtttgtaattgtccttttaacagcgatcgcgtattt
cgtctcgctcaggcgcaatcacgaatgaataacggtttggttgatgcgagtgattttgatgacgagcgtaatggctggcctgttgaacaagtctggaaagaa
atgcataagcttttgccattctcaccggattcagtcgtcactcatggtgatttctcacttgataaccttatttttgacgaggggaaattaataggttgtattgatg
ttggacgagtcggaatcgcagaccgataccaggatcttgccatcctatggaactgcctcggtgagttttctccttcattacagaaacggctttttcaaaaatat
ggtattgataatcctgatatgaataaattgcagtttcatttgatgctcgatgagtttttctaatcagaattggttaattggttgtaacactggcagagcattacg
ctgacttgacgggacggcacctgtagagaagagtccctgaatatcaaaatggtgggataaaaagctcaaaaaggaaagtaggctgtggttccctaggcaa
cagtcttccctaccccactggaaactaaaaaaacgagaaaagttcgcaccgaacatcaattgcataattttagccctaaaacataagctgaacgaaactgg
ttgtcttcccttcccaatccaggacaatctgagaatcccctgcaacattacttaacaaaaaagcaggaataaaattaacaagatgtaacagacataagtccc
atcaccgttgtataaagttaactgtgggattgcaaaagcattcaagcctaggcgctgagctgtttgagcatcccggtggcccttgtcgctgcctccgtgtttct
ccctggatttatttaggtaatatctctcataaatccccgggtagttaacgaaagttaatggagatcagtaacaataactctagggtcattactttggactccct
cagtttatccgggggaattgtgtttaagaaaatcccaactcataaagtcaagtaggagattaattcaatgaaaaattgcttgaaaatgaagaacctcctgcc
tgccttgaccatcacgatggcaatgagtgcagtgatggcgctggtcgtcacacctaacgcctatgcttctaaatgggatgagaaaatgactcctgagcaagt
ggaggccactctggataagaaatttgcagaaggaaactacagccctaaaggcgccgacagttgtctaatgtgccacaaaaagagcgagaaggtcatgga
tctcttcaagggtgttcatggcgccattgatagtagtaagagtcccatggctggtttgcagtgcgaagcctgccatgggccgctagggcaacacaacaaggg
gggcaacgagccgatgatcacgtttgggaaacagagtacactatctgcggacaagcaaaatagcgtctgtatgtcttgccaccaagacgataaacgtatg
agttggaatggagggcatcatgataatgctgacgtcgcctgcgcatcctgccaccaagttcatgtagcgaaggacccggtgttaagcaagaatacagaaat
ggaggtatgcacgtcctgtcataccaaacaaaaggctgacatgaacaagcgtagctcccatccgttaaagtgggcccagatgacatgcagcgactgccat
aacccccacgggtccatgaccgattctgacttgaacaagcccagtgtcaacgatacatgctatagttgtcatgcagagaaacgaggcccaaagctgtggga
acacgcgccggttacggaaaattgcgtgacgtgtcacaatcctcatggttccgtcaatgatggaatgctaaagacgcgcgcccctcaattatgccagcaatg
ccatgcgagcgatggtcacgcgagtaatgcgtacttaggtaatacgggactgggcagtaacgtgggggacaatgcgtttaccggaggacgaagctgcctc
aactgtcactctcaggtgcatggtagtaaccatccgagtggcaaactgttacagcga

**MtrA+(futA)**

aacgtatgagttggaatggagggcatcatgataatgctgacgtcgcctgcgcatcctgccaccaagttcatgtagcgaaggacccggtgttaagcaagaat
acagaaatggaggtatgcacgtcctgtcataccaaacaaaaggctgacatgaacaagcgtagctcccatccgttaaagtgggcccagatgacatgcagcg
actgccataacccccacgggtccatgaccgattctgacttgaacaagcccagtgtcaacgatacatgctatagttgtcatgcagagaaacgaggcccaaag
ctgtgggaacacgcgccggttacggaaaattgcgtgacgtgtcacaatcctcatggttccgtcaatgatggaatgctaaagacgcgcgcccctcaattatgc
cagcaatgccatgcgagcgatggtcacgcgagtaatgcgtacttaggtaatacgggactgggcagtaacgtgggggacaatgcgtttaccggaggacgaa
gctgcctcaactgtcactctcaggtgcatggtagtaaccatccgagtggcaaactgttacagcgaagatctgattataaagatcatgatggcgattataaag
atcatgatattgattataaagatgatgatgataaataataaggatccaacgctcggttgccgccgggcgtttttttattctgcaggagcagtagagcatacatc
tggaagcaaagccaggaaagcggcctatggagctgtgcggcagcgctcagtaggcaatttttcaaaatattgttaagccttttctgagcatggtatttttcat
ggtattaccaattagcaggaaaataagccattgaatataaaagataaaaatgtcttgtttacaatagagtgggggggtcagcctgccgccttgggccggg
tgatgtcgtacttgcccgccgcgaactcggttaccgtccagcccagcgcgaccagctccggcaacgcctcgcgcacccgctggcggcgcttgcgcatggtcg
aaccactggcctctgacggccagacatagccgcacaaggtatctatggaagccttgccggttttgccggggtcgatccagccacacagccgctggtgcagc
aggcgggcggtttcgctgtccagcgcccgcacctcgtccatgctgatgcgcacatgctggccgccacccatgacggcctgcgcgatcaaggggttcagggc
cacgtacaggcgcccgtccgcctcgtcgctggcgtactccgacagcagccgaaacccctgccgcttgcggccattctgggcgatgatggataccttccaaag
gcgctcgatgcagtcctgtatgtgcttgagcgccccaccactatcgacctctgccccgatttcctttgccagcgcccgatagctacctttgaccacatggcatt
cagcggtgacggcctcccacttgggttccaggaacagccggagctgccgtccgccttcggtcttgggttccgggccaagcactaggccattaggcccagcca
tggccaccagcccttgcaggatgcgcagatcatcagcgcccagcggctccgggccgctgaactcgatccgcttgccgtcgccgtagtcatacgtcacgtcca
gcttgctgcgcttgcgctcgccccgcttgagggcacggaacaggccgggggccagacagtgcgccgggtcgtgccggacgtggctgaggctgtgcttgttct
taggcttcaccacggggcaccccttgctcttgcgctgcctctccagcacggcgggcttgagcaccccgccgtcatgccgcctgaaccaccgatcagcgaac
ggtgcgccatagttggccttgctcacaccgaagcggacgaagaaccggcgctggtcgtcgtccacaccccattcctcggcctcggcgctggtcatgctcgac
aggtaggactgccagcggatgttatcgaccagtaccgagctgccccggctggcctgctgctggtcgcctgcgcccatcatggccgcgcccttgctggcatgg
tgcaggaacacgatagagcacccggtatcggcggcgatggcctccatgcgaccgatgacctgggccatggggccgctggcgttttcttcctcgatgtggaa
ccggcgcagcgtgtccagcaccatcaggcggcggccctcggcggcgcgcttgaggccgtcgaaccactccggggccatgatgttgggcaggctgccgatc

agcggctggatcagcaggccgtcagccacggcttgccgttcctcggcgctgaggtgcgccccaagggcgtgcaggcggtgatgaatggcggtgggcgggt
cttcggcgggcaggtagatcaccgggccggtgggcagttcgcccacctccagcagatccggcccgcctgcaatctgtgcggccagttgcagggccagcatg
gatttaccggcaccaccgggcgacaccagcgccccgaccgtaccggccaccatgttgggcaaaacgtagtccagcggtggcggcgctgctgcgaacgcct
ccagaatattgataggcttatgggtagccattgattgcctcctttgcaggcagttggtggttaggcgctggcggggtcactacccccgccctgcgccgctctg
agttcttccaggcactcgcgcagcgcctcgtattcgtcgtcggtcagccagaacttgcgctgacgcatccctttggccttcatgcgctcggcatatcgcgcttg
gcgtacagcgtcagggctggccagcaggtcgccggtctgcttgtccttttggtctttcatatcagtcaccgagaaacttgccggggccgaaaggcttgtcttc
gcggaacaaggacaaggtgcagccgtcaaggttaaggctggccatatcagcgactgaaaagcggccagcctcggccttgtttgacgtataaccaaagcca
ccgggcaaccaatagcccttgtcacttttgatcaggtagaccgaccctgaagcgctttttcgtattccataaaccccttctgtgcgtgagtactcatagtat
aacaggcgtgagtaccaacgcaagcactacatgctgaaatctggcccgcccctgtccatgcctcgctggcggggtgccggtgcccgtgccagctcggcccg
cgcaagctggacgctgggcagacccatgaccttgctgacggtgcgctcgatgtaatccgcttcgtggccgggcttgcgctctgccagcgctgggctggcctc
ggccatggccttgccgatttcctcggcactgcggccccggctggccagcttctgcgcggcgataaagtcgcacttgctgaggtcatcaccgaagcgcttgac
cagcccggccatctcgctgcggtactcgtccagcgccgtgcgccggtggcggctaagctgccgctcgggcagttcgaggctggccagcctgcgggccttctc
ctgctgccgctgggcctgctcgatctgctggccagcctgctgcaccagcgccgggccagcggtggcggtcttgcccttggattcacgcagcagcacccacgg
ctgataaccggcgcgggtggtgtgcttgtccttgcggttggtgaagcccgccaagcggccatagtggcggctgtcggcgctggccgggtcggcgtcgtactc
gctggccagcgtccgggcaatctgcccccgaagttcaccgcctgcggcgtcggccaccttgacccatgcctgatagttcttcgggctggtttccactaccagg
gcaggctcccggccctcggctttcatgtcatccaggtcaaactcgctgaggtcgtccaccagcaccagaccatgccgctcctgctcggcgggcctgatatac
acgtcattgccctgggcattcatccgcttgagccatggcgtgttctggagcacttcggcggctgaccattcccggttcatcatctggccggtggtggcgtccct
gacgccgatatcgaagcgctcacagcccatggccttgagctgtcggcctatggcctgcaaagtcctgtcgttcttcatcgggccaccaagcgcagccagatc
gagccgtcctcggttgtcagtggcgtcaggtcgagcaagagcaacgatgcgatcagcagcaccaccgtaggcatcatggaagccagcatcacggttagcc
atagcttccagtgccaccccgcgacgcgctccgggcgctctgcgcggcgctgctcacctcggcggctacctcccgcaactctttggccagctccacccatgc
cgcccctgtctggcgctgggctttcagccactccgccgcctgcgcctcgctggcctgctgggtctggctcatgacctgccgggcttcgtcggccagtgtcgcca
tgctctgggccagcggttcgatctgctccgctaactcgttgatgcctctggatttcttcactctgtcgattgcgttcatggtctattgcctcccggtattcctgtaa
gtcgatgatctgggcgttggcggtgtcgatgttcagggccacgtctgcccggtcggtgcggatgccccggccttccatctccaccacgttcggccccaggtga
acaccgggcaggcgctcgatgccctgcgcctcaagtgttctgtggtcaatgcgggcgtcgtggccagcccgctctaatgcccggttggcatggtcggcccat
gcctcgcgggtctgctcaagccatgccttgggcttgagcgcttcggtcttctgtgccccgcccttctccggggtcttgccgttgtaccgcttgaaccactgagcg
gcgggccgctcgatgccgtcattgatccgctcggagatcatcaggtggcagtgcgggttctcgccgccaccggcatggatggccagcgtatacggcaggcg
ctcggcaccggtcaggtgctgggcgaactcggacgccagcgccttctgctggtcgagggtcagctcgaccggcagggcaaattcgacctccttgaacagcc
gcccattggcgcgttcatacaggtcggcagcatcccagtagtcggcgggccgctcgacgaactccggcatgtgcccggattcggcgtgcaagacttcatcca
tgtcgcgggcatacttgccttcgcgctggatgtagtcggccttggccctggccgattggccgcccgacctgctgccggttttcgccgtaaggtgataaatcgcc
atgctgcctcgctgttgcttttgcttttcggctccatgcaatggccctcggagagcgcaccgcccgaagggtggccgttaggccagtttctcgaagagaaacc
ggtaagtgcgccctcccctacaaagtagggtcgggattgccgccgctgtgcctccatgatagcctacgagacagcacattaacaatggggtgtcaagatgg
ttaaggggagcaacaaggcggcggatcggctggccaagctcgaagaacaacgagcgcgaatcaatgccgaaattcagcgggtgcgggcaagggaacag
cagcaagagcgcaagaacgaaacaaggcgcaaggtgctggtgggggccatgattttggccaaggtgaacagcagcgagtggccggaggatcggctcat
ggcggcaatggatgcgtaccttgaacgcgaccacgaccgcgccttgttcggtctgccgccacgccagaaggatgagccgggctgaatgatcgaccgagac
aggccctgcggggctgcacacgcgcccccacccttcgggtagggggaaaggccgctaaagcggctaaaagcgctccagcgtatttctgcggggtttggtgt
ggggtttagcgggctttgcccgcctttcccctgccgcgcagcggtggggcggtgtgtagcctagcgcagcgaatagaccagctatccggcctctggccggg
catattgggcaagggcagcagcgccccacaagggcgctgataaccgcgcctagtggattattcttagataatcatggatggatttttccaacaccccgccag
cccccgcccctgctgggtttgcaggtttgggggcgtgacagttattgcaggggttcgtgacagttattgcagggggggcgtgacagttattgcaggggttcgtg
acagttagtacgggagtgacgggcactggctggcaatgtctagcaacggcaggcatttcggctgagggtaaaagaactttccgctaagcgatagactgtat
gtaaacacagtattgcaaggacgcggaacatgcctcatgtggcggccaggacggccagccgggatcgggatactggtcgttaccagagccaccgacccga
gcaaacccttctctatcagatcgttgacgagtattacccggcattcgctgcgcttatggcagagcagggaaaggaattgccgggctatgtgcaacgggaatt
tgaagaatttctccaatgcgggcggctggagcatggctttctacgggttcgctgcgagtcttgccacgccgagcacctggtcgctttcagctgtaatccgggc
agcgcaacggaacattcatcagtgtaaaaatggaatcaataaagccctgcgcagcgcgcagggtcagcctgaatacgcgtttaatgaccagcacagtcgt
gatggcaaggtcagaatagcgctgaggtctgcctcgtgaagaaggtgttgctgactcataccaggcctgaatcgccccatcatccagccagaaagtgaggg
agccacggttgatgagagctttgttgtaggtggaccagttggtgattttgaacttttgctttgccacggaacggtctgcgttgtcgggaagatgcgtgatctga
tccttcaactcagcaaaagttcgatttattcaacaaagccacgttgtgtctcaaaatctctgatgttacattgcacaagataaaaatatatcatcatgaacaa
taaaactgtctgcttacataaacagtaatacaaggggtgttatgagccatattcaacgggaaacgtcttgctcgaggccgcgattaaattccaacatggatg
ctgatttatatgggtataaatgggctcgcgataatgtcgggcaatcaggtgcgacaatctatcgattgtatgggaagcccgatgcgccagagttgtttctgaa
acatggcaaaggtagcgttgccaatgatgttacagatgagatggtcagactaaactggctgacggaatttatgcctcttccgaccatcaagcattttatccgt
actcctgatgatgcatggttactcaccactgcgatccccgggaaaacagcattccaggtattagaagaatatcctgattcaggtgaaaatattgttgatgcgc
tggcagtgttcctgcgccggttgcattcgattcctgtttgtaattgtccttttaacagcgatcgcgtatttcgtctcgctcaggcgcaatcacgaatgaataacg
gtttggttgatgcgagtgattttgatgacgagcgtaatggctggcctgttgaacaagtctggaaagaaatgcataagcttttgccattctcaccggattcagtc
gtcactcatggtgatttctcacttgataaccttatttttgacgaggggaaattaataggttgtattgatgttggacgagtcggaatcgcagaccgataccagg
atcttgccatcctatggaactgcctcggtgagttttctccttcattacagaaacggctttttcaaaaatatggtattgataatcctgatatgaataaattgcagt
ttcatttgatgctcgatgagtttttctaatcagaattggttaattggttgtaacactggcagagcattacgctgacttgacgggacggcacctgtagagaaga
gtccctgaatatcaaaatggtgggataaaaagctcaaaaaggaaagtaggctgtggttccctaggcaacagtcttccctaccccactggaaactaaaaaa
acgagaaaagttcgcaccgaacatcaattgcataattttagccctaaaacataagctgaacgaaactggttgtcttcccttcccaatccaggacaatctgag

aatcccctgcaacattacttaacaaaaaagcaggaataaaattaacaagatgtaacagacataagtcccatcaccgttgtataaagttaactgtgggattg
caaaagcattcaagcctaggcgctgagctgtttgagcatcccggtggcccttgtcgctgcctccgtgtttctccctggatttatttaggtaatatctctcataaa
tccccgggtagttaacgaaagttaatggagatcagtaacaataactctagggtcattactttggactccctcagtttatccgggggaattgtgtttaagaaaa
tcccaactcataaagtcaagtaggagattaattcaatgactacaaagattagtcgccgtacattctttgtaggaggaacggcgctaaccgcattagtagttg
caaatttgcctcgacgggcctctgcccagtcccggacatctaaatgggatgagaaaatgactcctgagcaagtggaggccactctggataagaaatttgca
gaaggaaactacagccctaaaggcgccgacagttgtctaatgtgccacaaaaagagcgagaaggtcatggatctcttcaagggtgttcatggcgccattg
atagtagtaagagtcccatggctggtttgcagtgcgaagcctgccatgggccgctagggcaacacaacaagggggcaacgagccgatgatcacgtttgg
gaaacagagtacactatctgcggacaagcaaaatagcgtctgtatgtcttgccaccaagacgata

**Empty vector**

agatctgattataaagatcatgatggcgattataaagatcatgatattgattataaagatgatgatgataaataataaggatccaacgctcggttgccgccg
ggcgtttttttattctgcaggagcagtagagcatacatctggaagcaaagccaggaaagcggcctatggagctgtgcggcagcgctcagtaggcaatttttca
aaatattgttaagccttttctgagcatggtatttttcatggtattaccaattagcaggaaaataagccattgaatataaaagataaaaatgtcttgtttacaat
agagtgggggggggtcagcctgccgccttgggccgggtgatgtcgtacttgcccgccgcgaactcggttaccgtccagcccagcgcgaccagctccggcaac
gcctcgcgcacccgctggcggcgcttgcgcatggtcgaaccactggcctctgacggccagacatagccgcacaaggtatctatggaagccttgccggttttg
ccggggtcgatccagccacacagccgctggtgcagcaggcgggcggtttcgctgtccagcgcccgcacctcgtccatgctgatgcgcacatgctggccgcc
acccatgacggcctgcgcgatcaaggggttcagggccacgtacaggcgcccgtccgcctcgtcgctggcgtactccgacagcagccgaaacccctgccgct
tgcggccattctgggcgatgatggataccttccaaaggcgctcgatgcagtcctgtatgtgcttgagcgccccaccactatcgacctctgccccgatttccttt
gccagcgcccgatagctacctttgaccacatggcattcagcggtgacggcctcccacttgggttccaggaacagccggagctgccgtccgccttcggtcttg
ggttccgggccaagcactaggccattaggcccagccatggccaccagcccttgcaggatgcgcagatcatcagcgcccagcggctccgggccgctgaactc
gatccgcttgccgtcgccgtagtcatacgtcacgtccagcttgctgcgcttgcgctcgccccgcttgagggcacggaacaggccgggggccagacagtgcgc
cgggtcgtgccggacgtggctgaggctgtgcttgttcttaggcttcaccacggggcaccccttgctcttgcgctgcctctccagcacggcgggcttgagcac
cccgccgtcatgccgcctgaaccaccgatcagcgaacggtgcgccatagttggccttgctcacaccgaagcggacgaagaaccggcgctggtcgtcgtcca
caccccattcctcggcctcggcgctggtcatgctcgacaggtaggactgccagcggatgttatcgaccagtaccgagctgccccggctggcctgctgctggtc
gcctgcgcccatcatggccgcgcccttgctggcatggtgcaggaacacgatagagcacccggtatcggcggcgatggcctccatgcgaccgatgacctggg
ccatggggccgctggcgttttcttcctcgatgtggaaccggcgcagcgtgtccagcaccatcaggcggcggccctcggcggcgcgcttgaggccgtcgaacc
actccggggccatgatgttgggcaggctgccgatcagcggctggatcagcaggccgtcagccacggcttgccgttcctcggcgctgaggtgcgccccaagg
gcgtgcaggcggtgatgaatggcggtgggcgggtcttcggcgggcaggtagatcaccgggccggtgggcagttcgcccacctccagcagatccggcccgc
ctgcaatctgtgcggccagttgcagggccagcatggatttaccggcaccaccgggcgacaccagcgccccgaccgtaccggccaccatgttgggcaaaac
gtagtccagcggtggcggcgctgctgcgaacgcctccagaatattgataggcttatgggtagccattgattgcctcctttgcaggcagttggtggttaggcgc
tggcggggtcactaccccgccctgcgccgctctgagttcttccaggcactcgcgcagcgcctcgtattcgtcgtcggtcagccagaacttgcgctgacgcat
ccctttggccttcatgcgctcggcatatcgcgcttggcgtacagcgtcagggctggccagcaggtcgccggtctgcttgtccttttggtctttcatatcagtcac
cgagaaacttgccggggccgaaaggcttgtcttcgcggaacaaggacaaggtgcagccgtcaaggttaaggctggccatatcagcgactgaaaagcggc
cagcctcggccttgtttgacgtataaccaaagccaccgggcaaccaatagcccttgtcacttttgatcaggtagaccgaccctgaagcgctttttttcgtattcc
ataaaaccccccttctgtgcgtgagtactcatagtataacaggcgtgagtaccaacgcaagcactacatgctgaaatctggcccgcccctgtccatgcctcgct
ggcggggtgccggtgcccgtgccagctcggcccgcgcaagctggacgctgggcagacccatgaccttgctgacggtgcgctcgatgtaatccgcttcgtgg
ccgggcttgcgctctgccagcgctgggctggcctcggccatggccttgccgatttcctcggcactgcggccccggctggccagcttctgcgcggcgataaagt
cgcacttgctgaggtcatcaccgaagcgcttgaccagcccggccatctcgctgcggtactcgtccagcgccgtgcgccggtggcggctaagctgccgctcgg
gcagttcgaggctggccagcctgcgggccttctcctgctgccgctgggcctgctcgatctgctggccagcctgctgcaccagcgccgggccagcggtggcgg
tcttgcccttggattcacgcagcagcacccacggctgataaccggcgcgggtggtgtgcttgtccttgcggttggtgaagcccgccaagcggccatagtggc
ggctgtcggcgctggccgggtcggcgtcgtactcgctggccagcgtccgggcaatctgcccccgaagttcaccgcctgcggcgtcggccaccttgacccatg
cctgatagttcttcgggctggtttccactaccagggcaggctcccggccctcggctttcatgtcatccaggtcaaactcgctgaggtcgtccaccagcaccag
accatgccgctcctgctcggcgggcctgatatacacgtcattgccctgggcattcatccgcttgagccatggcgtgttctggagcacttcggcggctgaccatt
cccggttcatcatctggccggtggtggcgtccctgacgccgatatcgaagcgctcacagcccatggccttgagctgtcggcctatggcctgcaaagtcctgtc
gttcttcatcgggccaccaagcgcagccagatcgagccgtcctcggttgtcagtggcgtcaggtcgagcaagagcaacgatgcgatcagcagcaccaccgt
aggcatcatggaagccagcatcacggttagccatagcttccagtgccacccccgcgacgcgctccgggcgctctgcgcggcgctgctcacctcggcggcta
cctcccgcaactctttggccagctccacccatgccgcccctgtctggcgctgggctttcagccactccgccgcctgcgcctcgctggcctgctgggtctggctc
atgacctgccgggcttcgtcggccagtgtcgccatgctctgggccagcggttcgatctgctccgctaactcgttgatgcctctggatttcttcactctgtcgatt
gcgttcatggtctattgcctcccggtattcctgtaagtcgatgatctgggcgttggcggtgtcgatgttcagggccacgtctgcccggtcggtgcggatgcccc
ggccttccatctccaccacgttcggccccaggtgaacaccgggcaggcgctcgatgccctgcgcctcaagtgttctgtggtcaatgcgggcgtcgtggccag
cccgctctaatgcccggttggcatggtcggcccatgcctcgcgggtctgctcaagccatgccttgggcttgagcgcttcggtcttctgtgccccgcccttctccg
gggtcttgccgttgtaccgcttgaaccactgagcggcgggccgctcgatgccgtcattgatccgctcggagatcatcaggtggcagtgcgggttctcgccgcc
accggcatggatggccagcgtatacggcaggcgctcggcaccggtcaggtgctgggcgaactcggacgccagcgccttctgctggtcgagggtcagctcg
accggcagggcaaattcgacctccttgaacagccgcccattggcgcgttcatacaggtcggcagcatcccagtagtcggcgggccgctcgacgaactccgg
catgtgcccggattcggcgtgcaagacttcatccatgtcgcgggcatacttgccttcgcgctggatgtagtcggccttggccctggccgattggccgcccgac
ctgctgccggttttcgccgtaaggtgataaatcgccatgctgcctcgctgttgcttttgcttttcggctccatgcaatggccctcggagagcgcaccgcccgaa
gggtggccgttaggccagtttctcgaagagaaaccggtaagtgcgccctcccctacaaagtagggtcgggattgccgccgctgtgcctccatgatagcctac

gagacagcacattaacaatggggtgtcaagatggttaaggggagcaacaaggcggcggatcggctggccaagctcgaagaacaacgagcgcgaatcaa
tgccgaaattcagcgggtgcgggcaagggaacagcagcaagagcgcaagaacgaaacaaggcgcaaggtgctggtgggggccatgattttggccaagg
tgaacagcagcgagtggccggaggatcggctcatggcggcaatggatgcgtaccttgaacgcgaccacgaccgcgccttgttcggtctgccgccacgccag
aaggatgagccgggctgaatgatcgaccgagacaggccctgcggggctgcacacgcgcccccacccttcgggtagggggaaaggccgctaaagcggcta
aaagcgctccagcgtatttctgcggggtttggtgtggggtttagcgggctttgcccgcctttcccccctgccgcgcagcggtggggcggtgtgtagcctagcgc
agcgaatagaccagctatccggcctctggccgggcatattgggcaagggcagcagcgccccacaagggcgctgataaccgcgcctagtggattattcttag
ataatcatggatggatttttccaacaccccgccagcccccgcccctgctgggtttgcaggtttgggggcgtgacagttattgcaggggttcgtgacagttattg
caggggggcgtgacagttattgcaggggttcgtgacagttagtacgggagtgacgggcactggctggcaatgtctagcaacggcaggcatttcggctgagg
gtaaaagaactttccgctaagcgatagactgtatgtaaacacagtattgcaaggacgcggaacatgcctcatgtggcggccaggacggccagccgggatc
gggatactggtcgttaccagagccaccgacccgagcaaacccttctctatcagatcgttgacgagtattacccggcattcgctgcgcttatggcagagcagg
gaaaggaattgccgggctatgtgcaacgggaatttgaagaatttctccaatgcgggcggctggagcatggctttctacgggttcgctgcgagtcttgccacg
ccgagcacctggtcgctttcagctgtaatccgggcagcgcaacggaacattcatcagtgtaaaaatggaatcaataaagccctgcgcagcgcgcagggtca
gcctgaatacgcgtttaatgaccagcacagtcgtgatggcaaggtcagaatagcgctgaggtctgcctcgtgaagaaggtgttgctgactcataccaggcct
gaatcgccccatcatccagccagaaagtgagggagccacggttgatgagagctttgttgtaggtggaccagttggtgattttgaacttttgctttgccacgga
acggtctgcgttgtcgggaagatgcgtgatctgatccttcaactcagcaaaagttcgatttattcaacaaagccacgttgtgtctcaaaatctctgatgttaca
ttgcacaagataaaaatatatcatcatgaacaataaaactgtctgcttacataaacagtaatacaaggggtgttatgagccatattcaacgggaaacgtctt
gctcgaggccgcgattaaattccaacatggatgctgatttatatgggtataaatgggctcgcgataatgtcgggcaatcaggtgcgacaatctatcgattgt
atgggaagcccgatgcgccagagttgtttctgaaacatggcaaaggtagcgttgccaatgatgttacagatgagatggtcagactaaactggctgacggaa
tttatgcctcttccgaccatcaagcattttatccgtactcctgatgatgcatggttactcaccactgcgatccccgggaaaacagcattccaggtattagaaga
atatcctgattcaggtgaaaatattgttgatgcgctggcagtgttcctgcgccggttgcattcgattcctgtttgtaattgtccttttaacagcgatcgcgtattt
cgtctcgctcaggcgcaatcacgaatgaataacggtttggttgatgcgagtgattttgatgacgagcgtaatggctggcctgttgaacaagtctggaaagaa
atgcataagcttttgccattctcaccggattcagtcgtcactcatggtgatttctcacttgataaccttatttttgacgaggggaaattaataggttgtattgatg
ttggacgagtcggaatcgcagaccgataccaggatcttgccatcctatggaactgcctcggtgagttttctccttcattacagaaacggctttttcaaaaatat
ggtattgataatcctgatatgaataaattgcagtttcatttgatgctcgatgagtttttctaatcagaattggttaattggttgtaacactggcagagcattacg
ctgacttgacgggacggcacctgtagagaagagtccctgaatatcaaaatggtgggataaaaagctcaaaaaggaaagtaggctgtggttccctaggcaa
cagtcttccctaccccactggaaactaaaaaaacgagaaaagttcgcaccgaacatcaattgcataattttagccctaaaacataagctgaacgaaactgg
ttgtcttcccttcccaatccaggacaatctgagaatcccctgcaacattacttaacaaaaaagcaggaataaaattaacaagatgtaacagacataagtccc
atcaccgttgtataaagttaactgtgggattgcaaaagcattcaagcctaggcgctgagctgtttgagcatcccggtggcccttgtcgctgcctccgtgtttct
ccctggatttatttaggtaatatctctcataaatccccgggtagttaacgaaagttaatggagatcagtaacaataactctagggtcattactttggactccct
cagtttatccgggggaattgtgtttaagaaaatcccaactcataaagtcaagtaggagattaattcaatgagaagagcgaattcggcgcgccggtaagatc
caaatcgatgaattgaccaagcactacggtatgaagccagaagactacactgctgtcagatgtggtatgaatgtcgccaagtacatcatcgaagataagat
tgatgctggtattggtatcgaatgtatgcaacaagtcgaattggaagagtacttggccaagcaaggcagaccagcttctgatgctaaaatgttgagaattga
caagttggcttgcttgggttgctgttgtttctgtaccgttctttacatctgcaacgatgaatttttgaagaaaaaccctgaaaaggtcagaaagttcttgaaag
ccatcaagaaggcaaccgactacgttctagccgaccctgtgaaggcttggaaagaatacatcgacttcaagcctcaattgaacagctcttca

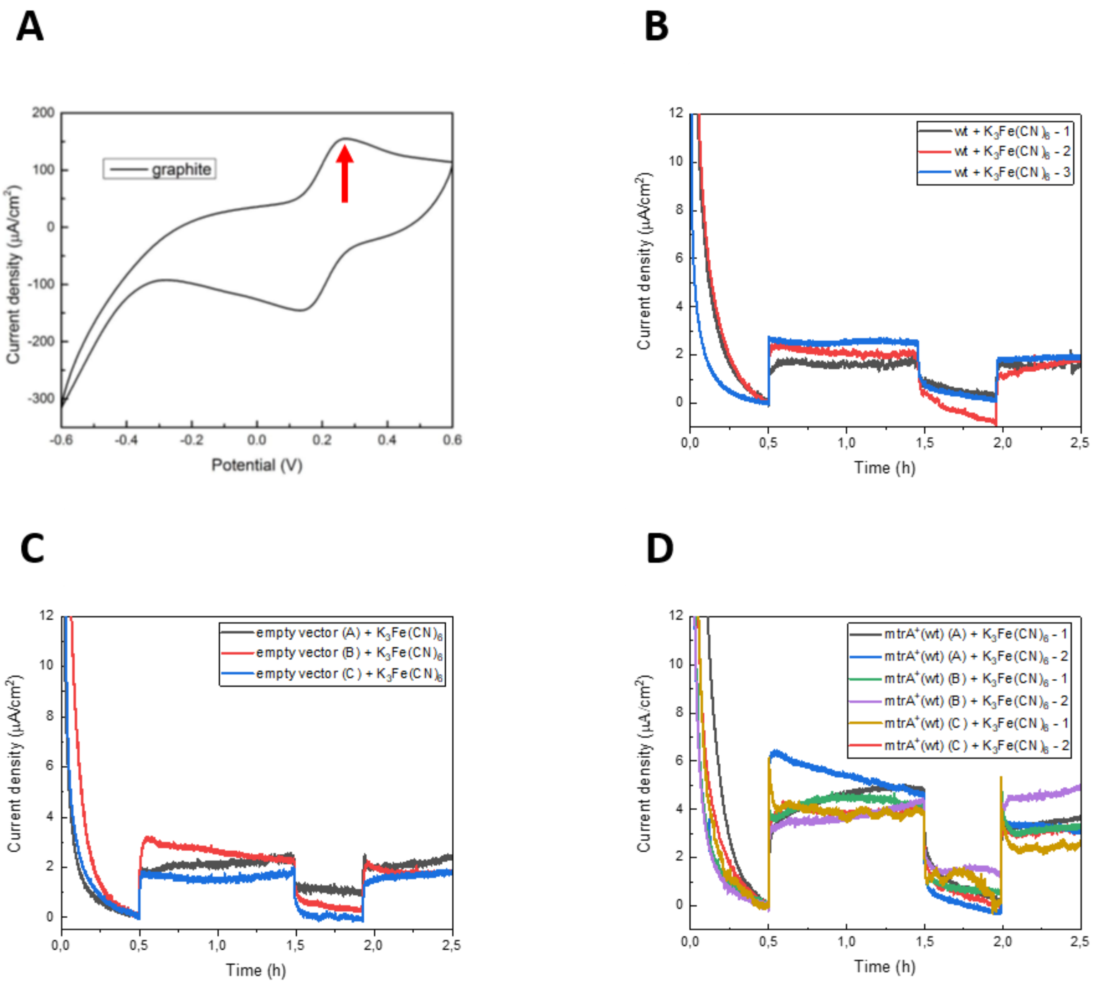


**Figure S2: (A)** Cyclic voltammograms of a 1 mM $K_4Fe(CN)_6$ aqueous solution performed to select the $[Fe(CN)_6]^{4-}$ oxidative potential (ca. 300 mV, red arrow). Individual chronoamperometry (CA) measurements showing the light response of wild-type *Synechocystis* sp. PCC 6803 **(B)** empty vector control **(C)** and mtrA+ expressing cells **(D)** on graphite electrodes. All the CAs lasted for 2.5 hours. The system was stabilized in the dark for 30 minutes, followed by 1 hour incubation under 500 µmol m-2s-1 of LED light illumination, 30 minutes of dark incubation, and another 30 minutes under the light.

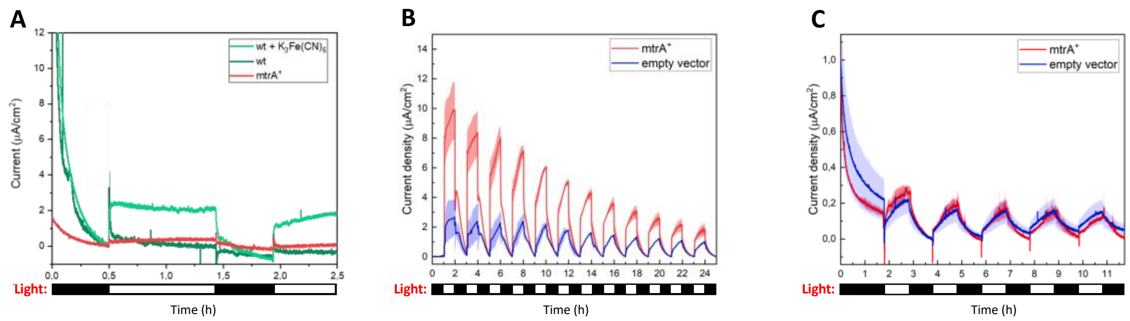


**Figure S3: (A)** Representative mediatorless light responses of wild-type *Synechocystis* sp. PCC 6803 (dark green) and MtrA-expressing cells (red) showing only a slight increase in current upon illumination compared to the one achieved by wt cells under $K_3Fe(CN)_6$-mediated conditions (light green). **(B)** Long term mediated and **(C)** unmediated CA measurements of mtrA$^+$ (red) and empty vector cells (blue). MtrA-expressing cells and empty vector cells exhibit similar currents under mediatorless conditions, because MtrA is not exposed on the outer membrane. All the CA measurements were taken using three technical replicates. The bold line corresponds to the mean value of the current density, and the shaded regions represent the standard deviation (SD).
